# Hemodynamic and Yap1/Wwtr1-dependent epithelial-endothelial signaling establish fenestrated endothelial identity and permeability in the choroid plexus

**DOI:** 10.64898/2026.11.21.768335

**Authors:** Jun Xiong Leong, Vani Thakur, Nathanael J. Lee, Ella N. Kazazic, Angelene A. Getz, Fatma Z. Bozdag, Ryota L. Matsuoka

## Abstract

The choroid plexuses (CPs) constitute the interface between cerebrospinal fluid (CSF) and circulating blood, mediating their molecular exchange. How the CPs establish permeable endothelial identities and properties distinct from neighboring barrier-forming endothelium remains unknown. By developing new genetic and functional screening platforms in zebrafish, we identified blood flow-responsive KLF2 orthologues, *klf2a* and *klf2b*, as critical regulators of CP endothelial identity and permeability. Subsequent investigations linked these findings to hemodynamic and mechanosensitive Yap1/Wwtr1 signaling requirements. Expression analyses showed Yap1/Wwtr1 in endothelial and epithelial CP compartments, while epistatic analyses suggested that Yap1/Wwtr1 activity counteracts reduced hemodynamic inputs in the endothelium. Conversely, Yap1/Wwtr1 activity regulates epithelial expression of the VEGFA orthologue *vegfab*, which signals through endothelial Vegfr2 in CP vascular specialization. These findings define hemodynamic and Yap1/Wwtr1-dependent epithelial-endothelial signaling as crucial determinants of CP vascular specialization and propose that systemic and organ-specific fluid mechanics intersect to shape tissue environments generating unique cellular properties.

## Introduction

During development, vascular networks expand to deliver nutrients, gases, and bioactive molecules throughout the body while undergoing regional specialization to meet the distinct physiological demands of individual organs. This tissue-specific organization generates heterogeneous vascular endothelial cell (EC) identities and functions within interconnected vascular networks (*1, 2*), with endothelial permeability and barrier properties representing prominent examples of vascular heterogeneity. Yet, how such heterogeneous EC identities and properties adapted to local tissue environments are established remains an important question in organogenesis and developmental biology.

The brain is a unique organ exhibiting striking intra-organ vascular heterogeneity in endothelial permeability and barrier properties (*3, 4*). In most regions of the brain, the vasculature acquires highly specialized barrier properties characterized by the blood-brain barrier (BBB) (*5, 6*). In contrast, the choroid plexuses (CPs) and circumventricular organs (CVOs) develop fenestrated, highly permeable capillaries that lack typical BBB properties (*3, 4*). These contrasting vascular functions support region-specific neural functions and brain homeostasis. Although extensive efforts have been made to study the mechanisms underlying BBB formation and integrity, much less is known about the developmental processes specifying fenestrated cerebrovascular identities and properties.

The CPs are evolutionarily conserved brain structures that produce cerebrospinal fluid (CSF), secrete growth factors into CSF to regulate brain development, control CSF homeostasis via molecular exchange between blood and CSF, form the blood-CSF barrier, and orchestrate neuroimmune surveillance (*7, 8*). These vital physiological functions are supported at least in part by their specialized fenestrated vasculature. PLVAP is an integral structural component of endothelial fenestrations and diaphragms and has been widely used to characterize permeable vascular beds across organs (*9, 10*). Within the central nervous system (CNS), immature cerebrovascular ECs exhibit higher PLVAP expression before acquiring mature BBB properties (*11*). Furthermore, PLVAP upregulation is observed following BBB disruption in various neurological conditions (*12*). Together, these findings have established PLVAP as a valuable marker for assessing vascular permeability states.

Previous studies have reported the role of Sonic hedgehog in regulating vascular outgrowth in the mouse hindbrain CP (*13*) and that VEGF-A and TGF-β signaling pathways contribute to the maintenance of fenestrated vascular properties in several cerebrovascular beds, including the adult CPs (*14, 15*). However, the molecular cues that establish endothelial fenestrations in developing CP vessels have yet to be identified. VEGF-A has been broadly recognized as a key regulator of endothelial fenestrations and permeability based on its demonstrated functions both *in vitro* and *in vivo* (*14–17*). Nevertheless, the upstream mechanisms that drive its expression in specific brain regions *in vivo* remain largely unclear. Hemodynamic forces are also well-established regulators of vascular remodeling, endothelial polarity, and vessel stability (*18, 19*). However, little is known about how systemic mechanical cues cooperate with organ-specific biochemical signals in vascular specialization.

Our recent studies have identified endothelial origins, angiogenic cues, and their cellular sources contributing to vascular patterning across multiple CVOs and CPs (*20–23*). However, these investigations primarily focused on defining the developmental mechanisms governing vascular patterning in these organs, leaving the following fundamental question unanswered: how are fenestrated cerebrovascular identities and properties specified during development? To date, the molecular and biophysical cues that establish permeable CP vasculature remain unknown across vertebrates. To address this knowledge gap, we investigated this understudied aspect of cerebrovascular biology.

Here, using the zebrafish hindbrain myelencephalic choroid plexus (mCP) as an *in vivo* model of highly permeable cerebrovasculature, we performed a CRISPR/Cas9-based genetic screen using F0 knockout larvae (crispants). Through this screen and subsequent analyses, we identified hemodynamic forces and associated molecular pathways as critical regulators of endothelial identity and permeability in the developing mCP. Further mechanistic analyses suggest a model in which systemic hemodynamic signaling and local epithelial-endothelial communication govern CP vascular specialization.

## Results

### Establishment of a new vascular permeability assay reveals cerebrovascular permeability heterogeneity and prominent leakage hotspots associated with the CPs and CVOs

Our previous work using conventional dextran tracers identified high vascular permeability in select cerebrovascular beds, including the diencephalic choroid plexus (dCP)-pineal gland interface (*20*). Compared with rapidly diffusing dextran tracers, we found that fluorescently labeled wheat germ agglutinin (WGA) at approximately 38 kDa, a lectin that binds to selective cell-surface glycoproteins (*24*), provided enhanced spatial visualization of extravasated tracers in both the dCP and mCP regions (Fig. 1a, b). We therefore established a WGA-based tracer assay as a valuable platform to examine and characterize developmental progression of regionally restricted high vascular permeability in these brain regions. Following intravascular injection of fluorescent WGA, we observed pronounced tracer extravasation within and around the dCP and mCP as early as 75 hours post-fertilization (hpf) (Fig. 1c-e). In contrast, substantially less extravasation was detected around neighboring BBB-type brain vessels at this stage, revealing notable regional heterogeneity in cerebrovascular permeability (Fig. 1f-h). Direct side-by-side comparisons of fluorescent WGA and a similarly sized dextran tracer (40 kDa) further showed that WGA provides clearer spatial visualization of accumulated tracer extravasation (Fig. 1c-f”).

**Fig. 1.**
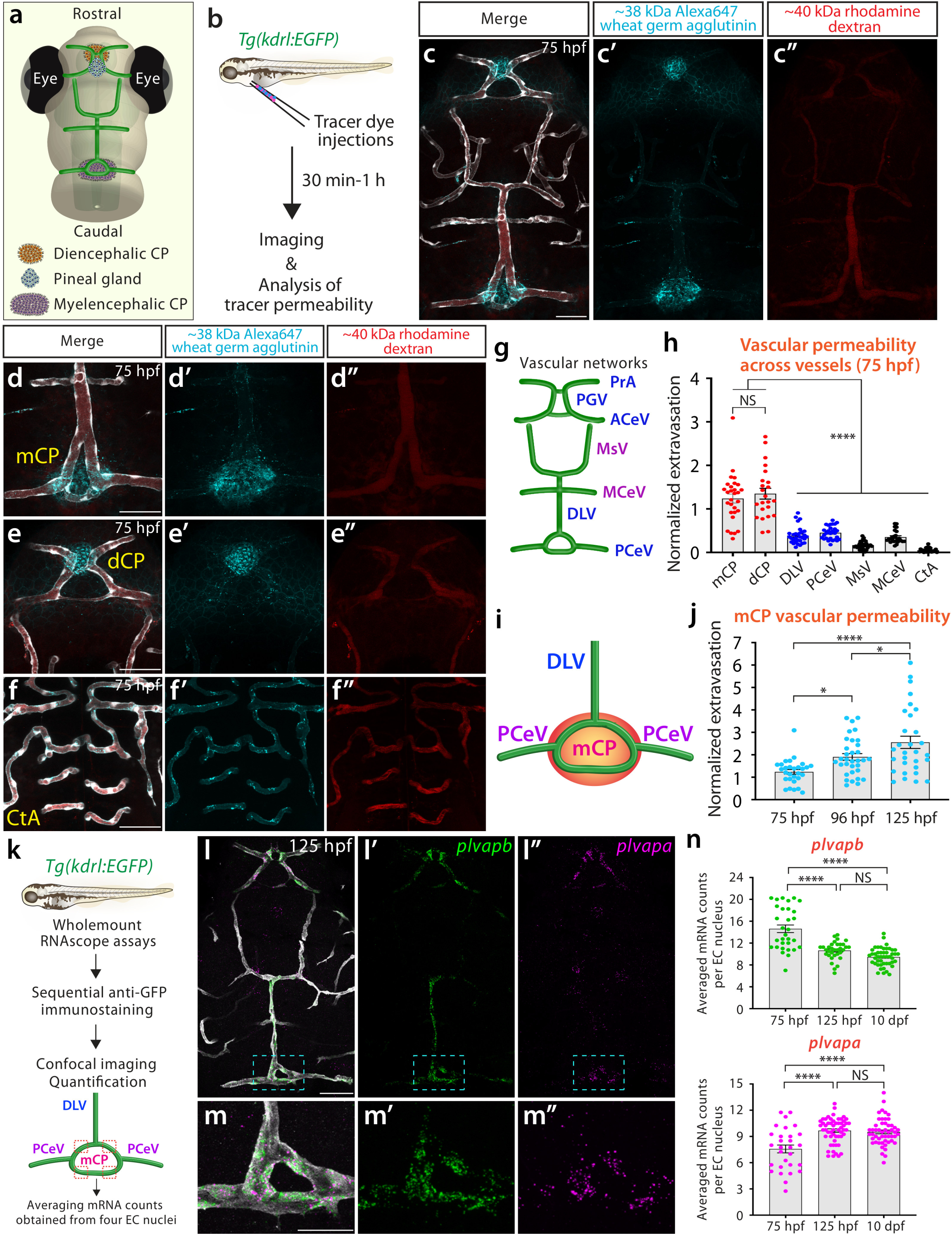
Cerebrovascular permeability heterogeneity and prominent leakage hotspots associated with the CPs and CVOs. (a) Schematic dorsal view of the zebrafish larval brain illustrating the diencephalic choroid plexus (dCP), pineal gland, and myelencephalic choroid plexus (mCP) and their associated vasculature. (b) Schematic of the vascular permeability assay. Fluorescent tracers were injected intravascularly into *Tg(kdrl:EGFP)* larvae, which were subsequently imaged 30 min-1 h after injection to assess tracer extravasation. (**c**-**f**) Representative dorsal views of 75 hpf larvae following co-injection of ∼38-kDa Alexa Fluor 647-conjugated wheat germ agglutinin (WGA; cyan; **c’**-**f’**) and ∼40-kDa rhodamine dextran (red; **c”**-**f”**). Merged images include *Tg(kdrl:EGFP)*-positive vasculature shown in gray. Prominent WGA extravasation is observed around the mCP (**d**) and dCP (**e**) vasculature, in contrast to the relatively impermeable BBB-type central arteries (CtAs; **f**). The WGA tracer enables clearer visualization of localized tracer extravasation in both CP regions compared with dextran of similar molecular size. (g) Schematic illustration of cranial vessels located near the dorsal brain surface and analyzed at 75 hpf. **PrA**: prosencephalic artery, **PGV**: pineal gland vessel, **ACeV**: anterior cerebral vein, **MsV**: mesencephalic vein, **MCeV**: middle cerebral vein, **DLV**: dorsal longitudinal vein, **PCeV**: posterior cerebral vein. (h) Quantification of normalized WGA extravasation across the indicated vascular beds at 75 hpf. (i) Schematic illustration of vessels contributing to the mCP vasculature and analyzed for vascular permeability. (j) Quantification of normalized WGA extravasation in the mCP vascular loop at 75, 96, and 125 hpf. (k) Experimental workflow for whole-mount RNAscope analysis of *plvapa* and *plvapb* expression in mCP endothelial cells (ECs) using the *Tg(kdrl:EGFP)* reporter. Transcript abundance was quantified in four ECs and averaged per sample. (**l**-**m”**) Representative dorsal views of *plvapb* (green) and *plvapa* (magenta) expression in the *Tg(kdrl:EGFP)* larval brain at 125 hpf. Magnified images of the mCP vasculature (boxed areas in **l**-**l”**) are shown in (**m**-**m”**). (n) Quantification of *plvapb* and *plvapa* mRNA counts in mCP ECs at 75 hpf, 125 hpf, and 10 dpf. Average mCP endothelial *plvapb* expression decreased from 75 to 125 hpf, whereas *plvapa* expression increased over the same period. Both expression levels remained stable from 125 hpf to 10 dpf. Scale bars: 50 µm in **c** for **c**-**c”**, **d** for **d**-**d”**, **e** for **e**-**e”**, **f** for **f**-**f”**, **l** for **l**-**l”**; 25 µm in **m** for **m**-**m”**.

We focused our subsequent analyses on the mCP due to its stereotypical vascular organization and tissue accessibility for quantitative imaging, and next examined developmental progression of vessel permeability in this vascular bed. Quantification of WGA extravasation revealed a progressive increase in vascular permeability from 75 to 125 hpf (Fig. 1i, j), indicating that high permeability of the mCP vasculature is established during early development and continues to increase through 125 hpf. To ask whether the developmental emergence of high vascular permeability correlates with an acquisition of fenestrated endothelial molecular identities, we performed whole-mount RNAscope analysis of the zebrafish PLVAP orthologues, *plvapa* and *plvapb* (*25*) (Fig. 1k). Using the pan-endothelial zebrafish reporter line, *Tg(kdrl:EGFP)*, both transcripts were prominently detected in vascular ECs of the dCP and mCP at 125 hpf (Fig. 1l-m”). Notably, *plvapa* expression exhibited a more spatially restricted distribution, whereas *plvapb* was expressed more broadly across vessels comprising these vascular beds, including the dorsal longitudinal vein (DLV) and bilateral posterior cerebral veins (PCeVs).

To characterize their developmental expression dynamics, we quantified both *plvapa* and *plvapb* transcript abundance in individual ECs by selecting two ECs from the rostral side and another two from the caudal side of the mCP vascular loop and averaged these four measurements to represent a biological sample (Fig. 1k). At 75 hpf, *plvapb* expression was relatively high with the greatest variability among the examined stages and subsequently decreased by 125 hpf, whereas *plvapa* expression progressively increased over the same developmental interval (Fig. 1n). Neither transcript exhibited significant changes in abundance per EC between 125 hpf and 10 days post fertilization (dpf) (Fig. 1n). These developmental expression dynamics define mCP endothelial *plvap* expression in relation to the development of high vascular permeability in the mCP. In particular, the developmental increase in *plvapa* expression from 75 to 125 hpf closely mirrored the increase in mCP vascular permeability observed during this period, indicating a temporal correlation between these changes. Together, these findings provide a foundation for utilizing spatially and temporally resolved, quantitative *plvapa*/*plvapb* expression analyses as molecular readouts of EC specialization in the permeable mCP vasculature.

### A CRISPR/Cas9-based F0 genetic screen identifies candidate regulators of robust *plvapa* expression in the mCP vasculature

To identify candidate regulators of mCP vascular specialization, we performed a CRISPR/Cas9-based F0 genetic screen using our newly established *plvapa/plvapb* expression platform. We narrowed down and selected candidate genes from several broad functional categories, including regulators of vascular permeability, transcription factors enriched in fenestrated endothelium, crucial genes in lymphatic and venous vascular biology, and regulators involved in inflammatory and immune signaling. To achieve efficient disruption of selected target genes, we co-injected synthetic CRISPR RNAs (crRNAs), trans-activating CRISPR RNA (tracrRNA), and Cas9 protein into one-cell-stage embryos using our previously established protocol employing clustered injections with three independent crRNAs per gene (*26, 27*).

For the purpose of facilitating this genetic screening and investigating potential functional redundancy among target genes, we strategically co-injected six crRNAs targeting two different genes to achieve simultaneous gene disruptions in some cases. The resulting F0 knockout larvae (crispants) were collected at 125 hpf for wholemount RNAscope analysis of *plvapa* and *plvapb* expression as molecular readouts of mCP endothelial states (Fig. 2a). In total, we screened 25 candidate genes across 20 genetic perturbation conditions (Fig. 2b). Quantification of transcript abundance identified a group of candidate genes whose disruption reduced *plvapa* expression compared with uninjected wild-type (WT) controls (Fig. 2b). In contrast, *plvapb* expression remained relatively stable across most examined conditions (Fig. 2b). Thus, this genetic screen successfully identified candidate regulators of robust *plvapa* expression in the mCP vasculature and suggested distinct regulatory requirements for *plvapa* and *plvapb* expression.

**Fig. 2.**
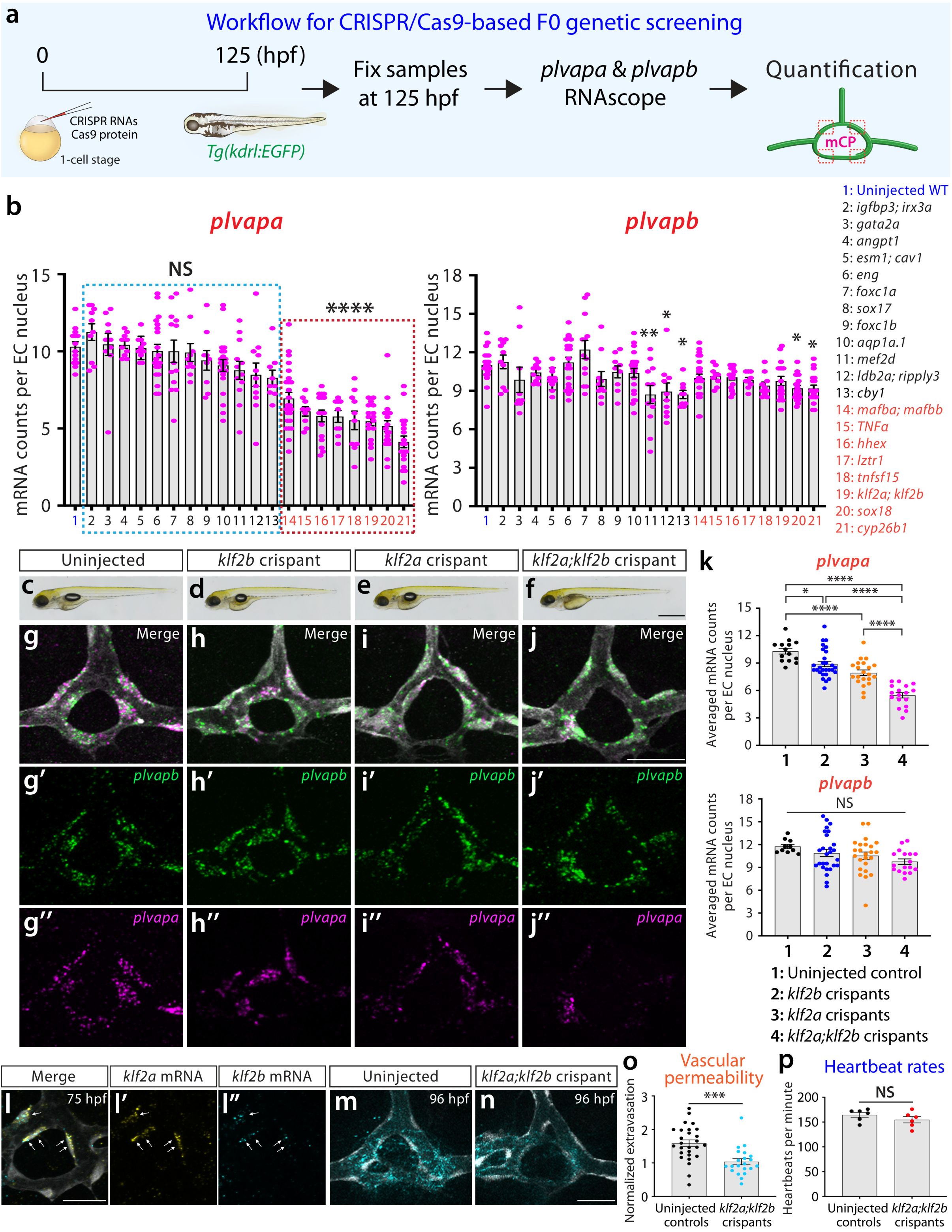
A genetic screen identifies Klf2 transcription factors as regulators of endothelial *plvapa* expression and permeability in the mCP. (a) Experimental workflow for targeted CRISPR/Cas9-based F0 genetic knockout (crispant) screening. One-cell-stage embryos were injected with gene-specific CRISPR RNAs and Cas9 protein, fixed at 125 hpf, and subjected to RNAscope analysis of *plvapa* and *plvapb* expression in mCP ECs. (b) Quantification of *plvapa* and *plvapb* mRNA abundance in mCP ECs at 125 hpf following CRISPR RNAs/Cas9 injections targeting the indicated candidate genes. Uninjected controls and targeted gene groups are numbered 1-21, as indicated. Combined targeting of *klf2a* and *klf2b* markedly reduced *plvapa* expression without significantly affecting *plvapb* expression (group 19). Statistical significance was determined by one-way ANOVA followed by Dunnett’s multiple comparisons test against uninjected controls. Asterisks above individual groups in the *plvapb* graph indicate statistically significant differences compared with uninjected controls. (**c**-**f**) Representative brightfield images of uninjected control, *klf2b* crispant, *klf2a* crispant, and *klf2a;klf2b* double-crispant larvae at 125 hpf. (**g**-**j”**) Representative dorsal views of mCP endothelial *plvapb* (green, **g’**-**j’**) and *plvapa* (magenta, **g”**-**j”**) expression in the indicated groups (merged images shown in **g**-**j**). While *plvapb* expression remained comparable across all conditions, *plvapa* expression was modestly reduced in *klf2b* and *klf2a* single crispants and markedly reduced in double crispants. (k) Quantification of *plvapa* and *plvapb* mRNA abundance in mCP ECs from uninjected controls, *klf2b*, *klf2a*, and *klf2a;klf2b* crispants. (**l–l”**) RNAscope analysis of *klf2a* (yellow, **l’**) and *klf2b* (cyan, **l”**) mRNA expression in the mCP vasculature at 75 hpf (merged image shown in **l**). Both *klf2a* and *klf2b* transcripts were detected within the same ECs (arrows) in this region. (**m**, **n**) Representative images of vascular dye leakage in the mCP of uninjected controls (**m**) and *klf2a;klf2b* double crispants (**n**) at 96 hpf, showing reduced WGA extravasation in the double crispants. (o) Quantification of normalized WGA extravasation in the mCP vasculature of uninjected controls and *klf2a;klf2b* double crispants at 96 hpf. (p) Quantification of heartbeat rates in uninjected controls and *klf2a;klf2b* double crispants at 125 hpf. Scale bars: 1 mm in **f** for **c**-**f**; 25 µm in **j** for **g**-**j”**, **l** for **l**-**l”**, **n** for **m**-**n**.

### Klf2 transcription factors regulate endothelial identity and permeability in the mCP vasculature

Among the identified candidates, we selected to focus on *klf2a* and *klf2b*, zebrafish orthologues of the mechanosensitive transcription factor KLF2, a key regulator of flow-dependent endothelial gene expression and vascular development (*28, 29*). Their established roles in endothelial responses to hemodynamic forces allowed us to investigate the understudied question of how biophysical cues and blood flow regulate CP vascular specialization. To examine their individual contributions to *plvapa* expression, we analyzed *klf2a* and *klf2b* single crispants in addition to *klf2a;klf2b* double crispants, which were originally identified as a positive hit in the screen (Fig. 2c-f). RNAscope analysis revealed significantly reduced *plvapa* expression in the mCP vasculature following disruption of either gene, with the most pronounced reduction observed in double crispants, as confirmed by quantitative analysis (Fig. 2k). By contrast, *plvapb* transcript abundance was not significantly altered across these conditions (Fig. 2k). Collectively, these findings indicate that both Klf2a and Klf2b contribute to robust *plvapa* expression in the mCP vasculature.

We next examined spatial expression patterns of *klf2a* and *klf2b* during mCP development. At 75 and 125 hpf, transcripts of both genes were detected within the same mCP ECs, supporting their potential roles in mediating endothelial mechanotransduction in these cells (Fig. 2l-l”; Fig. S1). To investigate whether disruption of *klf2a* and *klf2b* also affects mCP vascular permeability, we performed WGA-based tracer assays in *klf2a;klf2b* double crispants at 96 hpf. We chose this developmental stage because WT mCP vessels exhibit high vascular permeability with relatively less variability (Fig. 1j), while limited pigmentation at this stage permits reliable visualization and quantification of tracer extravasation in live larvae. Quantification revealed significantly reduced mCP WGA extravasation in double crispants compared with uninjected controls (Fig. 2m-o). Importantly, we measured heartbeat rates and observed no significant difference between the two groups (Fig. 2p), suggesting that the reduced vascular permeability was unlikely to be attributed to altered heartbeat rates, a physiological factor that can affect systemic blood circulation. Together, these genetic and functional findings identified Klf2a and Klf2b as crucial regulators of endothelial *plvapa* expression and permeability in the mCP. Notably, these subsequent analyses validated the effectiveness of our genetic screen and provided the first example of genes required for permeable mCP vascular properties.

### Hemodynamic forces as mechanical regulators of mCP endothelial identity and permeability

Given the established roles of endothelial Klf2 transcription factors in responding to hemodynamic forces (*28, 29*), we next asked whether blood flow contributes to permeable mCP endothelial specialization (Fig. 3a). Since endothelial *plvapa* expression and vascular permeability progressively increase in the mCP between 75 and 125 hpf (Fig. 1j, n), we sought to manipulate hemodynamic activity during this developmental period. To reduce cardiac activity and consequently alter blood flow, we adapted previously described pharmacological methods (*30–32*). Specifically, we treated larvae with the voltage-gated sodium channel blocker tricaine at two concentrations and exposure durations (either 150 mg/L for 48 h from 77 to 125 hpf or 450 mg/L for 24 h from 101 to 125 hpf), or with the L-type calcium channel inhibitor nifedipine at 30 μM for 24 h from 101 to 125 hpf (Fig. 3b).

**Fig. 3.**
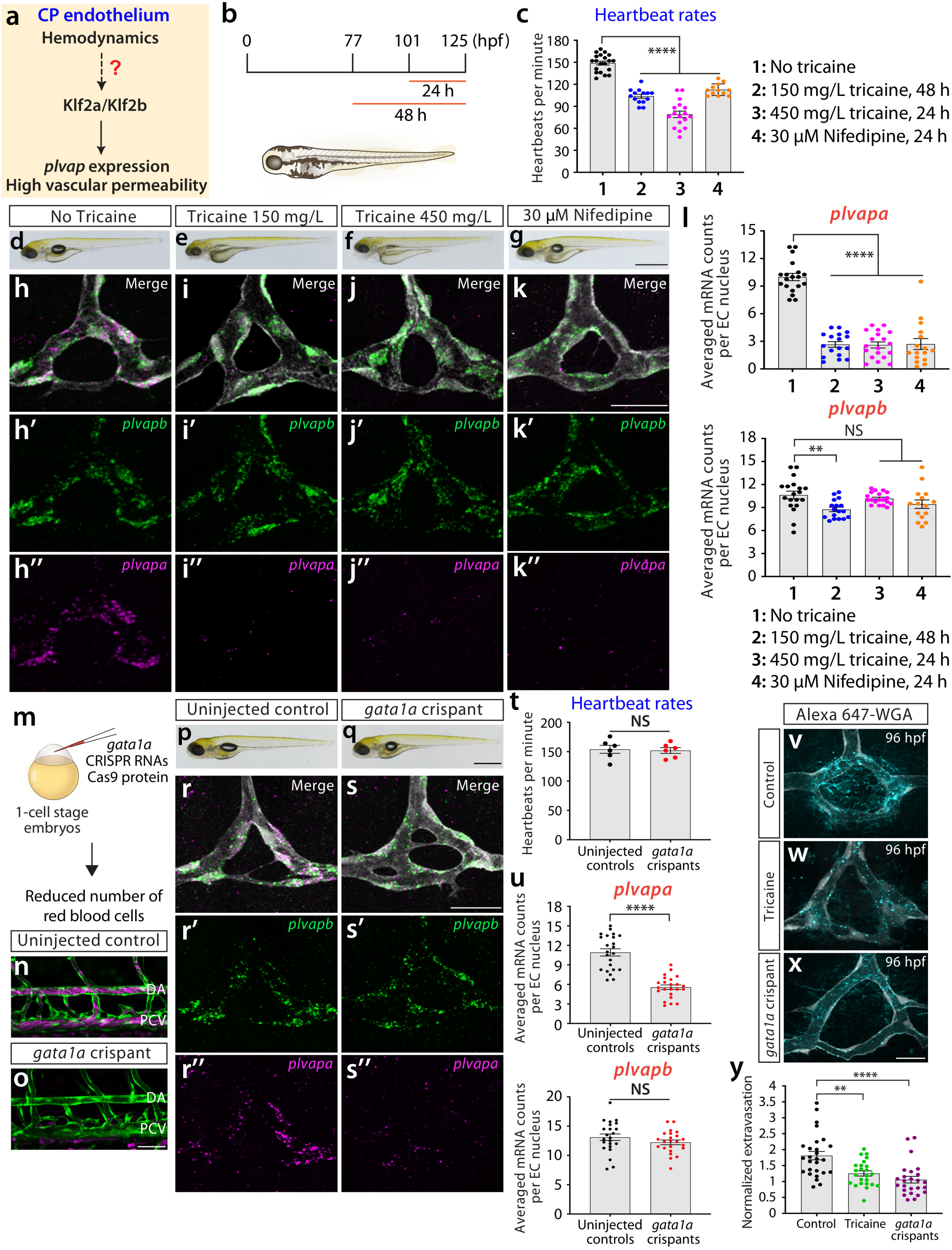
Hemodynamic activity is required for endothelial *plvapa* expression and vascular permeability in the mCP. (a) Schematic illustration of the central question addressed in subsequent experiments designed to investigate a potential role for hemodynamics in regulating *plvap* expression and vascular permeability in the CP endothelium. (**b**, **c**) Experimental workflow for pharmacological treatments (**b**) aimed at reducing cardiac activity and associated systemic circulation. Larvae were treated with 150 mg/L tricaine for 48 h, 450 mg/L tricaine for 24 h, or 30 µM nifedipine for 24 h before sample collection at 125 hpf. Treatment with the indicated tricaine or nifedipine resulted in markedly reduced heart rates compared to controls at 125 hpf (**c**). (**d**-**g**) Brightfield images of untreated (**d**), 150 mg/L tricaine-treated for 48 h (**e**), 450 mg/L tricaine-treated for 24 h (**f**), and 30 µM nifedipine-treated for 24 h (**g**) larvae at 125 hpf. (**h**-**k”**) Dorsal view images of *plvapb* (green, **h’**-**k’**) and *plvapa* (magenta, **h”**-**k”**) expression in the mCP of the indicated groups at 125 hpf (merged images shown in **h**-**k**). Pharmacological reduction of cardiac activity markedly decreased *plvapa* expression, whereas *plvapb* expression was largely preserved. (l) Quantification of *plvapa* and *plvapb* mRNA abundance in mCP ECs of untreated controls and larvae subjected to the indicated tricaine or nifedipine treatments at 125 hpf. (m) Experimental workflow for CRISPR/Cas9-mediated reduction of erythrocytes. One-cell-stage embryos were injected with *gata1a*-specific CRISPR RNAs and Cas9 protein to generate crispants. (**n**, **o**) Lateral view images of the trunk vasculature in uninjected controls (**n**) and *gata1a* crispants (**o**) at 125 hpf, showing a drastic reduction in circulating erythrocytes in the crispants. DA: dorsal aorta; PCV: posterior cardinal vein. (**p**, **q**) Brightfield images of uninjected control (**p**) and *gata1a* crispant (**q**) larvae at 125 hpf. (**r**-**s”**) Dorsal view images of *plvapb* (green, **r’**-**s’**) and *plvapa* (magenta, **r”**-**s”**) expression in the mCP of uninjected controls (**r**-**r”**) and *gata1a* crispants (**s**-**s”**) at 125 hpf (merged images shown in **r**, **s**). While *plvapb* expression remained comparable to controls, *plvapa* expression was markedly reduced in *gata1a* crispants. (t) Quantification of heartbeat rates in uninjected controls and *gata1a* crispants at 125 hpf. (u) Quantification of *plvapa* and *plvapb* mRNA abundance in mCP ECs of uninjected controls and *gata1a* crispants at 125 hpf. (**v**-**x**) Dorsal view images of Alexa Fluor 647-WGA extravasation in the mCP of untreated control (**v**), tricaine-treated (**w**), and *gata1a* crispant (**x**) larvae at 96 hpf, showing reduced vascular dye leakage in the tricaine-treated and *gata1a* crispant larvae. (**y**) Quantification of normalized WGA extravasation in the mCP vasculature of untreated controls, tricaine-treated larvae, and *gata1a* crispants at 96 hpf. Scale bars: 25 µm in **k** for **h**-**k”**, **s** for **r**-**s”**; 50 µm in **o** for **n**, **o**; 20 µm in **x** for **v**-**x**; 1 mm in **g** for **d**-**g**, **q** for **p**, **q**.

Each treatment condition led to significantly reduced heart rates at 125 hpf, with the magnitude of reduction varying across conditions (Fig. 3c). We then examined *plvapa* and *plvapb* expression in mCP ECs at 125 hpf and found that both tricaine and nifedipine treatments led to dramatically reduced *plvapa* expression, whereas *plvapb* expression was comparatively less affected (Fig. 3d-l). Since blood flow-dependent mechanical forces activate endothelial mechanosensitive channels (*33–35*), we also examined the effects of the mechanosensitive ion channel inhibitor GsMTx4 (*36*) at 300 nM for 24 or 48 h, using the same exposure durations as in the tricaine experiments. Both treatment conditions significantly reduced *plvapa*, but not *plvapb*, expression (Fig. S2). However, crispants targeting *piezo1*, which encodes a major endothelial mechanosensitive ion channel (*33–35*), did not exhibit a comparable phenotype (Fig. S2), indicating that mechanosensitive ion channels other than Piezo1 or their cooperative functions may contribute to the regulation of endothelial *plvapa* expression.

Since circulating red blood cells contribute to blood viscosity and thereby affect hemodynamic activity (*37*), we next generated crispants targeting *gata1a*, a transcription factor critical for red blood cell differentiation during hematopoiesis (*38*) (Fig. 3m). As previously demonstrated using antisense morpholinos targeting this gene (*38*), *gata1a* crispants exhibited a substantial reduction in circulating red blood cells at 125 hpf (Fig. 3n, o). Importantly, we observed that heart rates in these crispants were comparable to those in uninjected controls (Fig. 3t). Thus, unlike our earlier pharmacological approaches that involved reduced cardiac activity, this approach of *gata1a* disruption was expected to perturb hemodynamic forces without directly suppressing heartbeats. Despite their preserved cardiac activity, *gata1a* crispants showed markedly reduced *plvapa* expression in mCP ECs, whereas *plvapb* expression remained unchanged at 125 hpf (Fig. 3p-s”, u). Combined with the pharmacological results, these complementary findings suggest that the developmental expression of *plvapa* depends on hemodynamic signaling and red blood cell-mediated mechanical forces.

We next asked whether reduced hemodynamic conditions also affect functional permeability of the mCP vasculature. To this end, we performed WGA-based tracer assays and found markedly reduced tracer extravasation in the mCP of *gata1a* crispants compared with uninjected controls (Fig. 3v, x, y). A similar reduction in tracer extravasation was observed following tricaine treatment (Fig. 3w, y), demonstrating that distinct perturbations of hemodynamic function converge on reduced mCP vascular permeability. Collectively, these findings identify hemodynamics as an important mechanical regulator of both the molecular and functional properties of the mCP vasculature.

### Vegfa signaling as a biochemical regulator of mCP endothelial identity and permeability

To gain a comparative perspective on hemodynamic contributions to mCP vascular properties, we next investigated growth factor signaling that may regulate this process. Given the previous work reporting a link between Klf2a and Vegf signaling (*30*), we sought to determine the role of Vegf signaling in this context. This pathway selection was also motivated by our recent studies identifying the crucial role of this signaling family in regulating the patterning of mCP vasculature (*20–22*). We thus focused our analysis on examining its specific role in regulating mCP vascular specialization (Fig. 4a).

**Fig. 4.**
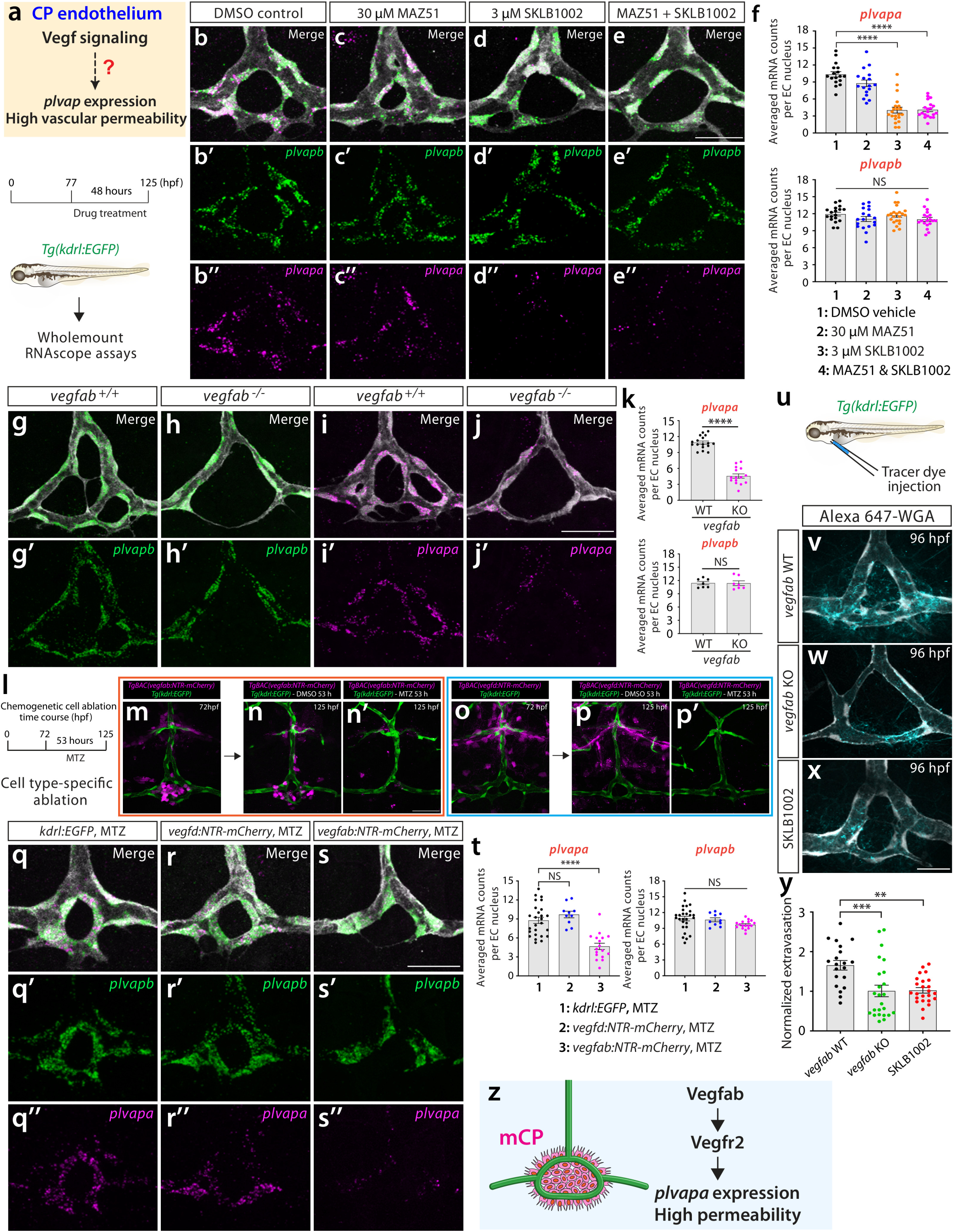
Vegfa-Vegfr2 signaling regulates endothelial *plvapa* expression and vascular permeability in the mCP. (**a**) Schematic illustration of the central question addressed in subsequent experiments designed to investigate a potential role for Vegf signaling in regulating *plvap* expression and vascular permeability in the CP endothelium. The lower section illustrates the experimental workflow for pharmacological inhibition of Vegf receptor signaling using the *Tg(kdrl:EGFP)* line. Larvae were treated with 30 µM MAZ51, 3 µM SKLB1002, or their combination for 48 h between 77 and 125 hpf before sample collection and subsequent whole-mount RNAscope analysis of *plvapa* and *plvapb* expression at 125 hpf. (**b**-**e”**) Dorsal view images of *plvapb* (green, **b’**-**e’**) and *plvapa* (magenta, **b”**-**e”**) expression in the mCP of DMSO-treated control (**b**-**b”**), MAZ51-treated (**c**-**c”**), SKLB1002-treated (**d**-**d”**), and combined MAZ51/SKLB1002-treated (**e**-**e”**) larvae at 125 hpf (merged images shown in **b**-**e**). While *plvapb* expression remained preserved across all conditions, *plvapa* expression was substantially reduced in SKLB1002-treated and combined MAZ51/SKLB1002-treated larvae. (**f**) Quantification of *plvapa* and *plvapb* mRNA abundance in mCP ECs of DMSO-treated controls and larvae subjected to the indicated pharmacological treatments at 125 hpf. (**g**-**j’**) Dorsal view images of *plvapb* (green, **g’**, **h’**) and *plvapa* (magenta, **i’**, **j’**) expression in the mCP of *vegfab* wild-type (WT) and homozygous mutant (KO) larvae at 125 hpf (merged images shown in **g**-**j**). Genetic loss of *vegfab* markedly reduced *plvapa* expression without affecting *plvapb* expression. (k) Quantification of *plvapa* and *plvapb* mRNA abundance in mCP ECs of *vegfab* WT and KO larvae at 125 hpf. (l) Experimental workflow for chemogenetic cell ablation using the nitroreductase (NTR)-metronidazole (MTZ) system. Larvae carrying the indicated *TgBAC(vegfab:gal4ff)* or *TgBAC(vegfd:gal4ff)* transgenes and *Tg(UAS:NTR-mCherry)*, abbreviated as *TgBAC(vegfab:NTR-mCherry)* and *TgBAC(vegfd:NTR-mCherry)*, respectively, were treated with MTZ from 72 to 125 hpf to induce cell-type-specific ablation. (**m**-**p’**) Representative images of *TgBAC(vegfab:NTR-mCherry)* (**m**-**n’**) and *TgBAC(vegfd:NTR-mCherry)* (**o**-**p’**) larvae carrying the *Tg(kdrl:EGFP)* transgene before (**m**, **o**) and after MTZ treatment (**n’**, **p’**), with DMSO-treated larvae serving as controls (**n**, **p**). Images were acquired at 72 hpf (**m, o**) and 125 hpf (**n, n’, p, p’**) to assess the effects of chemogenetic cell ablation. (**q**-**s”**) Dorsal view images of *plvapb* (green, **q’**-**s’**) and *plvapa* (magenta, **q”**-**s”**) expression in the mCP of MTZ-treated control (**q**-**q”**), *TgBAC(vegfd:NTR-mCherry)* (**r**-**r”**), and *TgBAC(vegfab:NTR-mCherry)* (**s**-**s”**) larvae at 125 hpf (merged images shown in **q**-**s**). While ablation of *vegfd*-expressing cells or MTZ treatment in controls did not alter *plvapa* expression, ablation of *vegfab*-expressing cells markedly reduced *plvapa* expression. *plvapb* expression remained comparable across these conditions. (t) Quantification of *plvapa* and *plvapb* mRNA abundance in mCP ECs following the indicated chemogenetic cell ablation and non-ablation treatments at 125 hpf. (u) Experimental workflow for vascular permeability analysis following intravascular injection of Alexa Fluor 647-WGA tracer into *Tg(kdrl:EGFP)* larvae. (**v**-**x**) Dorsal view images of Alexa Fluor 647-WGA extravasation in the mCP vasculature of *vegfab* WT (**v**), *vegfab* KO (**w**), and larvae treated with 3 µM SKLB1002 for 24 h (**x**) at 96 hpf. WGA extravasation was significantly reduced in SKLB1002-treated and *vegfab* KO larvae. (y) Quantification of normalized WGA extravasation in the mCP vasculature of *vegfab* WT, *vegfab* KO, and SKLB1002-treated larvae at 96 hpf. (z) Schematic diagram illustrating a critical role for Vegfab-Vegfr2 signaling in regulating endothelial *plvapa* expression and high vascular permeability in the mCP. Scale bars: 25 µm in **e** for **b**-**e”**, **j** for **g**-**j’**, **s** for **q**-**s”**; 50 µm in **n’** for **m**-**p’**; 20 µm in **x** for **v**-**x**.

To this end, we first treated larvae from 77 to 125 hpf for 48 h with the Vegfr2-specific inhibitor SKLB1002, the Vegfr3-specific inhibitor MAZ51, or both compounds, and conducted a quantitative RNAscope analysis of *plvapa* and *plvapb* expression (Fig. 4a). We found that SKLB1002 treatments led to a pronounced reduction in *plvapa* expression, whereas MAZ51 treatment had little detectable effect (Fig. 4b-d”, f). Combined treatment with SKLB1002 and MAZ51 did not further enhance *plvapa* reduction compared with SKLB1002 alone (Fig. 4d-f). In contrast, *plvapb* expression remained unchanged across all treatment conditions (Fig. 4f), suggesting a selective requirement of Vegfr2 signaling in regulating endothelial *plvapa* expression in the mCP. In support of these pharmacological observations, RNAscope analysis revealed mCP endothelial expression of the two zebrafish orthologues of VEGFR2, *kdr* and *kdrl* (*39*), at 125 hpf (Fig. S3).

To determine whether this effect reflected a specific requirement for Vegfa signaling, we next investigated how genetic *vegfa* loss contributes to this process. Zebrafish possess two Vegfa paralogs, Vegfaa and Vegfab (*40*), and our previous studies identified the importance of Vegfab in governing mCP vascular formation (*21, 22*). Thus, we examined *plvapa* and *plvapb* expression at 125 hpf in *vegfab* homozygous mutants and their WT siblings. Since approximately half of *vegfab* homozygous mutants exhibit defective DLV and/or PCeV formation, we restricted our analysis to mutants that successfully formed the mCP vasculature. We found that these *vegfab* mutants exhibited a pronounced reduction in *plvapa*, but not *plvapb*, expression in a manner closely resembling the effects of SKLB1002 treatment (Fig. 4g-k). Similarly, reducing Vegfa bioavailability via heatshock-induced overexpression of the soluble form of Vegfr1 (sFlt1), a decoy receptor for Vegfa (*41*), resulted in a selective reduction in *plvapa*, but not *plvapb*, expression in mCP ECs (Fig. S4). Together, these findings suggest that Vegfab-Vegfr2 signaling plays a crucial role in regulating *plvapa* expression in the mCP vasculature.

We next used a cell ablation strategy to selectively eliminate sources of Vegfab. In our previous study using the *vegfab* BAC transgenic reporter, we identified several cellular sources of *vegfab* near the mCP, including mCP epithelial cells and meningeal fibroblast-like cells (*21*). Since our study also indicated preferential *vegfd* BAC transgenic reporter expression in meningeal fibroblast-like cells, we sought to ablate these cells for comparison and to identify more precise cellular sources contributing to *plvapa* expression in the mCP vasculature. To this end, we induced cell ablation using the established nitroreductase (NTR)-metronidazole (MTZ) system (*42*). Specifically, we used larvae carrying either the *TgBAC(vegfab:gal4ff)* or *TgBAC(vegfd:gal4ff)* reporter together with the *Tg(UAS:NTR-mCherry)* transgene by administering the prodrug MTZ from 72 to 125 hpf (Fig. 4l). These treatments effectively eliminated the targeted Vegfab- or Vegfd-expressing cells (Fig. 4m-p’). We observed that ablation of *vegfab*-expressing cells markedly reduced *plvapa* expression in mCP endothelial cells, whereas ablation of *vegfd*-expressing cells produced no effect (Fig. 4q-t). Neither cell ablation affected mCP endothelial *plvapb* expression (Fig. 4q-t). These cell ablation experiments suggest a model in which mCP epithelial-derived local Vegfab signaling regulates *plvapa* expression in mCP endothelium via Vegfr2.

Finally, we conducted WGA tracer analysis and observed reduced mCP tracer extravasation in *vegfab* mutants compared with WT siblings (Fig. 4u-w). Pharmacological inhibition of Vegfr2 signaling with SKLB1002 similarly reduced tracer extravasation (Fig. 4x). Thus, both genetic and pharmacological perturbations of Vegfa signaling led to reduced mCP vascular permeability (Fig. 4y). Collectively, these findings identify Vegfab-Vegfr2 signaling as an important biochemical regulator of both the molecular and functional properties of the mCP vasculature (Fig. 4z), with mCP epithelial cells representing a potential local source of Vegfab.

### Yap1/Wwtr1 activity is required for mCP endothelial identity and permeability

Previous studies have demonstrated that Yes-associated protein 1 (Yap1) and WW-domain-containing transcription regulator 1 (Wwtr1) serve as transcriptional coactivators that respond to diverse biochemical and mechanical signals in the endothelium to control vascular functions (*29, 43, 44*). Given the identified requirements of hemodynamics and Vegfa signaling in mCP vascular specialization, we next asked whether Yap1/Wwtr1 activity contributes to the regulation of mCP endothelial properties and potentially integrates these distinct regulatory inputs (Fig. 5a).

**Fig. 5.**
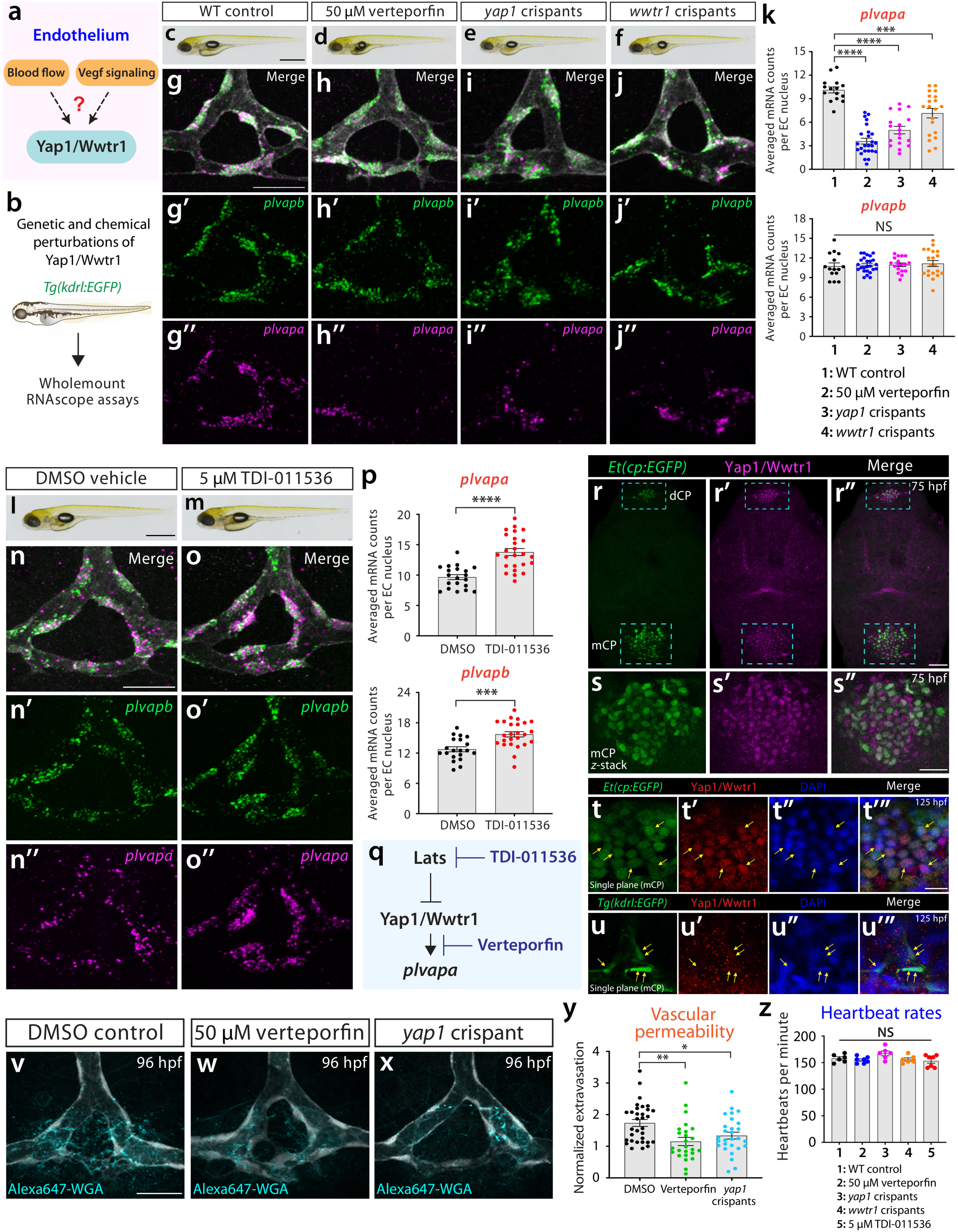
Yap1/Wwtr1 activity promotes endothelial *plvapa* expression and vascular permeability in the mCP. (a) Schematic illustration of the central question addressed in subsequent experiments designed to investigate a role for Yap1/Wwtr1 in mediating endothelial responses to blood flow and Vegf signaling. (b) Experimental design and workflow for testing genetic and pharmacological perturbations of Yap1/Wwtr1 signaling. *Tg(kdrl:EGFP)* larvae were used for the indicated perturbations and subsequent whole-mount RNAscope analysis of *plvapa* and *plvapb* expression in mCP ECs. (**c**-**f**) Brightfield images of WT control (**c**), 50 µM verteporfin-treated for 48 h (**d**), *yap1* crispant (**e**), and *wwtr1* crispant (**f**) larvae at 125 hpf. (**g**-**j”**) Dorsal view images of *plvapb* (green, **g’**-**j’**) and *plvapa* (magenta, **g”**-**j”**) expression in the mCP of the indicated groups at 125 hpf (merged images shown in **g**-**j**). *plvapa* expression was markedly reduced following verteporfin treatment and in *yap1* and *wwtr1* crispants, whereas *plvapb* expression remained preserved. (**k**) Quantification of *plvapa* and *plvapb* mRNA abundance in mCP ECs of WT controls, verteporfin-treated larvae, *yap1* crispants, and *wwtr1* crispants at 125 hpf. (**l**, **m**) Brightfield images of 125 hpf larvae after treatment with DMSO (**l**) or 5 µM TDI-011536 (**m**) for 48 h between 77 and 125 hpf. (**n**-**o”**) Dorsal view images of *plvapb* (green, **n’**, **o’**) and *plvapa* (magenta, **n”**, **o”**) expression in the mCP of DMSO-treated (**n**-**n”**) and TDI-011536-treated (**o**-**o”**) larvae at 125 hpf (merged images shown in **n, o**). TDI-011536 treatment significantly increased both *plvapa* and *plvapb* expression, with a more pronounced effect on *plvapa* expression. (p) Quantification of *plvapa* and *plvapb* mRNA abundance in mCP ECs of DMSO-treated and TDI-011536-treated larvae at 125 hpf. (q) Schematic illustration of pharmacological modulation of Yap1/Wwtr1 activity through LATS inhibition by TDI-011536 and disruption of Yap1/Wwtr1-dependent transcription by verteporfin. (**r**-**s”**) Dorsal view images of Yap1/Wwtr1 immunostaining (magenta, **r’**, **s’**) in the *Et(cp:EGFP)* (green, **r**, **s**) larval head at 75 hpf (merged images shown in **r”**, **s”**). The anterior and posterior boxed regions indicate the dCP and mCP, respectively. Prominent Yap1/Wwtr1 immunoreactivity was detected in both the dCP and mCP epithelium. Magnified *z*-stack images of the mCP region are shown in (**s**-**s”**). (**t**-**t’”**) Single-plane images of Yap1/Wwtr1 immunostaining (red, **t’**) and nuclear counterstaining with DAPI (blue, **t”**) in the *Et(cp:*EGFP*)^+^* larval mCP epithelium (green, **t**) at 125 hpf (merged image shown in **t”’**). Nuclear Yap1/Wwtr1 immunoreactivity was detected in the mCP epithelium. Arrows point to selected epithelial cells with clear nuclear Yap1/Wwtr1 immunoreactivity. (**u**-**u’”**) Single-plane images of Yap1/Wwtr1 immunostaining (red, **u’**) and nuclear counterstaining with DAPI (blue, **u”**) in the *Tg(kdrl:*EGFP*)^+^*larval mCP ECs (green, **u**) at 125 hpf (merged image shown in **u”’**). Nuclear Yap1/Wwtr1 immunoreactivity was detected in the mCP ECs (arrows). (**v**-**x**) Dorsal view images of Alexa Fluor 647-WGA extravasation in the mCP vasculature of DMSO-treated (**v**), 50 µM verteporfin-treated (**w**), and *yap1* crispant (**x**) larvae at 96 hpf, showing reduced vascular dye leakage in verteporfin-treated and *yap1* crispant larvae. (y) Quantification of normalized WGA extravasation in the mCP vasculature of DMSO-treated controls, verteporfin-treated larvae, and *yap1* crispants at 96 hpf. (z) Quantification of heartbeat rates in the indicated experimental groups. Scale bars: 25 µm in **g** for **g**-**j”**, **n** for **n**-**o”**, **s”** for **s**-**s”**; 50 µm in **r”** for **r**-**r”**; 10 µm in **t”’** for **t**-**t”’**, **u”’** for **u**-**u”’**; 20 µm in **v** for **v**-**x**; 1 mm in **c** for **c**-**f**, **l** for **l**, **m**.

To pursue this possibility, we first pharmacologically inhibited Yap1/Wwtr1 activity using verteporfin, a small-molecule inhibitor of YAP/WWTR1-dependent transcriptional activity (*45*), and examined *plvapa* and *plvapb* expression in mCP endothelial cells at 125 hpf (Fig. 5b-d, g-h”). Verteporfin treatment from 77 to 125 hpf for 48 h caused a pronounced reduction in *plvapa* expression, whereas *plvapb* expression remained unchanged (Fig. 5g-h”, k). This selective reduction in *plvapa* expression closely resembled the molecular phenotypes observed under perturbed hemodynamic forces or Vegfa signaling in the preceding experiments.

We next employed a genetic approach to investigate contributions of Yap1 and Wwtr1 to this process. To this end, we disrupted *yap1* or *wwtr1* using our established CRISPR/Cas9 injection strategy (*26, 27*) (Fig. 5e, f). We observed that both *yap1* and *wwtr1* crispants displayed reduced mCP endothelial *plvapa* expression, whereas *plvapb* expression remained unchanged (Fig. 5i-k). Together, these complementary pharmacological and genetic results support a role for Yap1/Wwtr1 signaling in regulating *plvapa* expression in the mCP vasculature.

Previous studies reported that large tumor suppressor kinases (LATS) phosphorylate YAP1/WWTR1 to restrict their nuclear localization and transcriptional activities (*46, 47*) and that LATS kinase inhibition increases YAP1/WWTR1 transcriptional activities (*48, 49*) (Fig. 5q). Thus, we next asked whether pharmacological activation of Yap1/Wwtr1 signaling alters *plvapa* and *plvapb* expression in mCP ECs. To this end, we treated larvae with TDI-011536, a small-molecule inhibitor of LATS kinases (*49*), from 77 to 125 hpf for 48 h to pharmacologically promote Yap1/Wwtr1 signaling and examined *plvapa* and *plvapb* expression at 125 hpf. We found that TDI-011536 treatment markedly increased *plvapa* expression compared with DMSO-treated controls (Fig. 5l-p). Interestingly, *plvapb* expression was also significantly increased, although the magnitude of this increase was less pronounced than that of *plvapa* (Fig. 5l-p). Together with our loss-of-function findings, these results support a role for Yap1/Wwtr1 signaling in regulating mCP endothelial *plvapa* expression and suggest that pharmacological LATS inhibition enhances both *plvapa* and *plvapb* expression in the mCP vasculature.

We next examined cellular localization of Yap1/Wwtr1 by performing immunostaining using an antibody against human YAP1/WWTR1 in the developing mCP. This antibody has been validated to recognize both human YAP1 and WWTR1 and used previously in zebrafish tissues, where it was reported to predominantly detect Wwtr1 (*50, 51*). Immunoreactivity with this antibody was prominently detected in both the dCP and mCP epithelium at 75 hpf (Fig. 5r-s”), with evident nuclear immunoreactivity in mCP epithelial cells at 125 hpf (Fig. 5t-t”’). We also observed fainter, yet detectable, nuclear signals in mCP vascular ECs at 125 hpf (Fig. 5u-u”’). These observations suggest that Yap1/Wwtr1 signaling is active in both mCP epithelial and endothelial populations during mCP vascular specialization, providing further support for their involvement in regulating this process.

Finally, we asked whether Yap1/Wwtr1 inhibition affects functional permeability of the mCP vasculature through WGA tracer assays. These analyses showed markedly reduced tracer extravasation in the mCP following verteporfin treatment and in *yap1* crispants compared with controls (Fig. 5v-y). Thus, pharmacological and genetic perturbations of Yap1/Wwtr1 signaling reduced both endothelial *plvapa* expression and vascular permeability in the mCP, identifying Yap1/Wwtr1 signaling as a potential transcriptional regulator of the molecular and functional properties of the mCP vasculature.

Because hemodynamic forces also regulate mCP endothelial specialization, we next examined whether these Yap1/Wwtr1 perturbations affected cardiac function. We found that neither verteporfin nor TDI-011536 treatment significantly altered heartbeat rates and that *yap1* and *wwtr1* crispants also showed no detectable changes compared with controls (Fig. 5z). Thus, the effects of Yap1/Wwtr1 perturbations on mCP endothelial *plvapa* expression and permeability are unlikely to be attributed to altered heartbeat rates, further supporting a role for Yap1/Wwtr1 signaling itself in these processes.

### Epistatic analysis of Yap1/Wwtr1 signaling in hemodynamic and Vegfa-dependent regulation of mCP vascular specialization

Previous studies showed that endothelial YAP1/WWTR1 signaling responds to blood flow-associated mechanical stimuli (*52–54*). Given our findings that both reduced hemodynamics and Yap1/Wwtr1 signaling decreased mCP endothelial *plvapa* expression, we hypothesized that Yap1/Wwtr1 signaling may act downstream of hemodynamics to regulate mCP endothelial specialization (Fig. 6a). To test this possibility, we combined *gata1a* disruption, which led to red blood cell depletion, with TDI-011536-mediated Lats inhibition and examined *plvapa* and *plvapb* expression at 125 hpf.

**Fig. 6.**
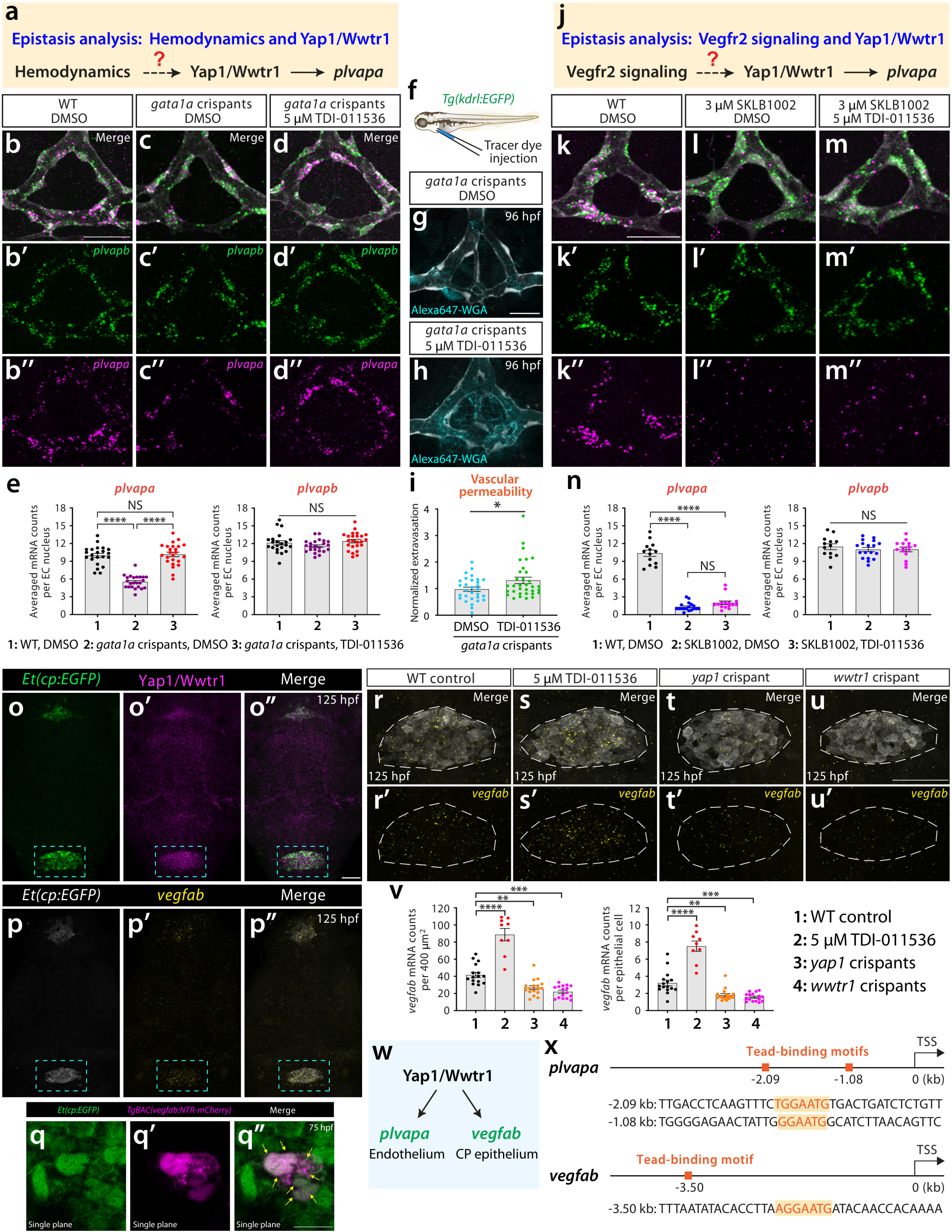
Yap1/Wwtr1 regulation of endothelial *plvapa* and epithelial *vegfab* expression in the mCP. (a) Schematic illustration of the central question addressed in (**b**-**e**) to investigate an epistatic relationship between hemodynamics and Yap1/Wwtr1 in mCP endothelial *plvapa* regulation. (**b**-**d”**) Dorsal view images of *plvapb* (green, **b’**-**d’**) and *plvapa* (magenta, **b”**-**d”**) expression in the mCP of DMSO-treated WT (**b**-**b”**), DMSO-treated *gata1a* crispant (**c**-**c”**), and 5 µM TDI-011536-treated *gata1a* crispant (**d**-**d”**) larvae at 125 hpf (merged images shown in **b**-**d**). Treatments were performed for 48 h between 77 and 125 hpf. TDI-011536 treatments rescued the reduced *plvapa* expression in *gata1a* crispants, whereas *plvapb* expression remained preserved. (e) Quantification of *plvapa* and *plvapb* mRNA abundance in mCP ECs under the indicated conditions. (f) Experimental workflow for vascular permeability analysis following intravascular injection of Alexa Fluor 647-WGA tracer into *Tg(kdrl:EGFP)* larvae. (**g**, **h**) Dorsal view images of the mCP vasculature of *gata1a* crispants treated with DMSO (**g**) or 5 µM TDI-011536 (**h**) for 24 h prior to Alexa Fluor 647-WGA injections at 96 hpf. TDI-011536 treatments significantly increased WGA extravasation in *gata1a* crispants. (i) Quantification of normalized WGA extravasation in the mCP vasculature of *gata1a* crispants at 96 hpf under the indicated treatments. (j) Schematic illustration of the central question addressed in (**k**-**n**) to investigate an epistatic relationship between Vegfr2 and Yap1/Wwtr1 in mCP endothelial *plvapa* regulation. (**k**-**m”**) Dorsal view images of *plvapb* (green, **k’**-**m’**) and *plvapa* (magenta, **k”**-**m”**) expression in the mCP of DMSO-treated controls (**k**-**k”**), 3 µM SKLB1002-treated larvae (**l**-**l”**), and larvae treated with 3 µM SKLB1002 and 5 µM TDI-011536 (**m**-**m”**) at 125 hpf (merged images shown in **k**-**m**). Treatments were performed for 48 h between 77 and 125 hpf. TDI-011536 treatments failed to rescue the reduced *plvapa* expression induced by SKLB1002. (**n**) Quantification of *plvapa* and *plvapb* mRNA abundance in mCP ECs under the indicated conditions. (**o**-**o”**) Dorsal view images of Yap1/Wwtr1 immunostaining (magenta, **o’**) in the *Et(cp:EGFP)* (green, **o**) larval head at 125 hpf (merged images shown in **o”**). Dashed boxes indicate the mCP region. (**p**-**p”**) Dorsal view images of *vegfab* mRNA expression (yellow, **p’**) in the *Et(cp:EGFP)* (gray, **p**) larval head at 125 hpf (merged image shown in **p”**). Dashed boxes indicate the mCP region. (**q**-**q”**) Dorsal view single-plane images of *Et(cp:EGFP)* (green, **q**) and *TgBAC(vegfab:NTR-mCherry)* (magenta, **q’**) expression in the mCP epithelium at 75 hpf (merged image shown in **q”**). Arrows point to double-positive cells showing *vegfab* BAC reporter expression in the CP epithelium. (**r**-**u’**) Dorsal view images of *vegfab* mRNA expression (yellow, **r’**-**u’**) in the *Et(cp:*EGFP)-positive mCP epithelium of WT (**r**, **r’**), 5 µM TDI-011536-treated (**s**, **s’**), *yap1* crispant (**t**, **t’**), and *wwtr1* crispant (**u**, **u’**) larvae at 125 hpf (merged images shown in **r**-**u**). TDI-011536 treatment for 48 h increased *vegfab* expression, whereas *yap1* and *wwtr1* disruption reduced its expression. Dashed outlines indicate the mCP region analyzed. (v) Quantification of *vegfab* mRNA abundance in mCP epithelial cells under the indicated conditions at 125 hpf. Graphs show mRNA counts per 400 µm^2^ area (left) and per mCP epithelial cell (right). (w) Schematic illustration summarizing Yap1/Wwtr1-dependent regulation of endothelial *plvapa* expression and epithelial *vegfab* expression in the mCP. (x) Schematic representation of predicted Tead-binding motifs within the 5 kb upstream regions of the zebrafish *plvapa* and *vegfab* genomic loci. Motifs were identified at approximately -2.09 and -1.08 kb upstream of the *plvapa* transcription start site (TSS) and approximately -3.50 kb upstream of the *vegfab* TSS. Predicted motifs are highlighted in orange within the corresponding nucleotide sequences. Scale bars: 25 µm in **b** for **b**-**d”**, **k** for **k**-**m”**; 20 µm in **g** for **g**, **h**; 50 µm in **o”** for **o**-**p”**, **u** for **r**-**u’**; 10 µm in **q”** for **q**-**q”**.

Consistent with our earlier findings, *gata1a* crispants treated with DMSO vehicle between 77 and 125 hpf for 48 h exhibited significantly reduced *plvapa* expression compared with controls (Fig. 6b-c”, e). Interestingly, TDI-011536 treatment during the same time window substantially increased *plvapa* expression in *gata1a* crispants, restoring it to levels comparable to those of controls (Fig. 6d-e). In contrast, *plvapb* expression remained unchanged across these experimental conditions (Fig. 6e). These findings suggest that enhancing Yap1/Wwtr1 activity can counteract the effects of reduced hemodynamic signaling and support a model in which Yap1/Wwtr1 signaling acts downstream of hemodynamics to regulate mCP endothelial *plvapa* expression.

We next examined whether this molecular recovery involved functional restoration of vascular permeability (Fig. 6f). TDI-011536 treatment alone did not significantly increase vascular permeability in WT larvae (Fig. S5), despite the increases in mCP endothelial *plvapa* and *plvapb* expression observed earlier. In contrast, TDI-011536 treatment significantly increased mCP vascular permeability in *gata1a* crispants, although permeability appeared to remain slightly below the levels observed in WT larvae (Fig. 6g-i; Fig. S5), suggesting a partial functional rescue. Together, these epistatic analyses demonstrate that enhancing Yap1/Wwtr1 activity counteracts the effects of reduced hemodynamic activity by restoring mCP endothelial *plvapa* expression and partially rescuing vascular permeability.

Since previous work suggested that Vegfa signaling also regulates Yap1/Wwtr1 activity in the endothelium (*44*), we asked whether a similar regulatory relationship exists between Yap1/Wwtr1 and Vegfa signaling (Fig. 6j). To test this idea, we treated larvae with the Vegfr2 inhibitor SKLB1002, either alone or in combination with TDI-011536 between 77 and 125 hpf for 48 h, and examined endothelial *plvapa* and *plvapb* expression at 125 hpf. Consistent with our previous results, SKLB1002 treatment alone led to a pronounced reduction in *plvapa* expression without significantly affecting *plvapb* expression (Fig. 6k-l”, n). However, TDI-011536 treatment failed to restore mCP endothelial *plvapa* expression under Vegfr2 inhibition (Fig. 6m-m”, n). This finding is in stark contrast to the ability of TDI-011536 to counteract the effects of reduced hemodynamic input. These findings imply distinct regulatory relationships between Yap1/Wwtr1 and hemodynamics or Vegfa signaling.

### Yap1/Wwtr1 activity regulates epithelial *vegfab* expression in the mCP

Epistasis analyses indicate that Yap1/Wwtr1 may function downstream of hemodynamic signaling in regulating mCP vascular specialization, whereas its relationship with Vegfa-Vegfr2 signaling appears to be clearly distinct. The inability of Lats inhibition to rescue mCP endothelial *plvapa* expression during Vegfr2 inhibition led us to hypothesize that Yap1/Wwtr1 instead act upstream of Vegfa signaling. Previous studies showed that YAP1 can promote VEGFA transcription in different cellular contexts (*55, 56*), providing further support for this hypothesis. Intriguingly, using the CP epithelial-specific reporter *Et(cp:EGFP)*, we found similar spatial distributions of Yap1/Wwtr1 immunoreactivity and *vegfab* expression within the mCP epithelium (Fig. 6o-p”). We further confirmed that *vegfab* BAC reporter expression overlaps with mCP epithelial cells labeled by the *Et(cp:EGFP)* reporter (Fig. 6q-q”). These observations are consistent with previous findings of constitutive VEGF-A expression in the adult mouse CP epithelium (*15*), indicating conserved CP epithelial VEGF-A expression across vertebrates and developmental stages.

The similar epithelial distributions of Yap1/Wwtr1 and *vegfab* led us to hypothesize that Yap1/Wwtr1 signaling regulates mCP epithelial *vegfab* expression through its transcriptional activity. To investigate this possibility, we performed RNAscope analysis and compared *vegfab* expression in mCP epithelial cells following pharmacological and genetic perturbations of Yap1/Wwtr1 signaling. We found that TDI-011536 treatment between 77 and 125 hpf for 48 h markedly increased epithelial *vegfab* expression compared with WT controls (Fig. 6r-s’, v). Conversely, genetic disruption of either *yap1* or *wwtr1* significantly reduced epithelial *vegfab* expression (Fig. 6t-v). These effects were consistent when *vegfab* transcripts were quantified either within a defined epithelial area or per epithelial cell. Collectively, these findings identify an unexpected role for Yap1/Wwtr1 signaling in regulating *vegfab* expression in the mCP epithelium.

Finally, to explore potential transcriptional mechanisms underlying Yap1/Wwtr1 regulation of endothelial *plvapa* and epithelial *vegfab* in the mCP (Fig. 6w), we examined genomic sequences upstream of their annotated transcription start sites (TSSs). To this end, we focused our analysis on finding consensus binding motifs for Tead transcription factors, well-established binding partners of Yap1/Wwtr1 (*57, 58*). Sequence analysis identified two candidate Tead binding motifs (5′-TGGAATG-3′ and 5′-GGAATG-3′) located approximately 2.09 and 1.08 kb, respectively, upstream of the *plvapa* TSS (Fig. 6x). Similarly, we identified one candidate Tead binding motif (5′-AGGAATG-3′) located approximately 3.50 kb upstream of the *vegfab* TSS (Fig. 6x). These sequence analyses indicate that Yap1/Wwtr1 may directly control transcription of both genes through Tead-dependent mechanisms.

## Discussion

Across vertebrates, the CPs constitute a unique physiological interface between two distinct fluid systems: CSF and circulating blood (*59, 60*). The conserved anatomical organization raises the intriguing possibility that distinct fluid mechanics shape the establishment of specialized functions in this organ. Here, our study identifies, for the first time in vertebrates, molecular and physiological cues that govern the development of fenestrated endothelial identity and high vascular permeability in the CP. By establishing genetic and functional screening platforms targeting the zebrafish hindbrain mCP, we demonstrate that Klf2a/Klf2b, hemodynamics, Vegfa signaling, and Yap1/Wwtr1 are critical regulators of endothelial *plvapa* expression and permeability in the developing mCP. Our subsequent investigations further illuminate mechanistic links among these pathways, providing a novel conceptual framework for understanding how systemic and organ-specific fluid mechanics may intersect to generate specialized CP vascular properties (Fig. 7).

**Fig. 7.**
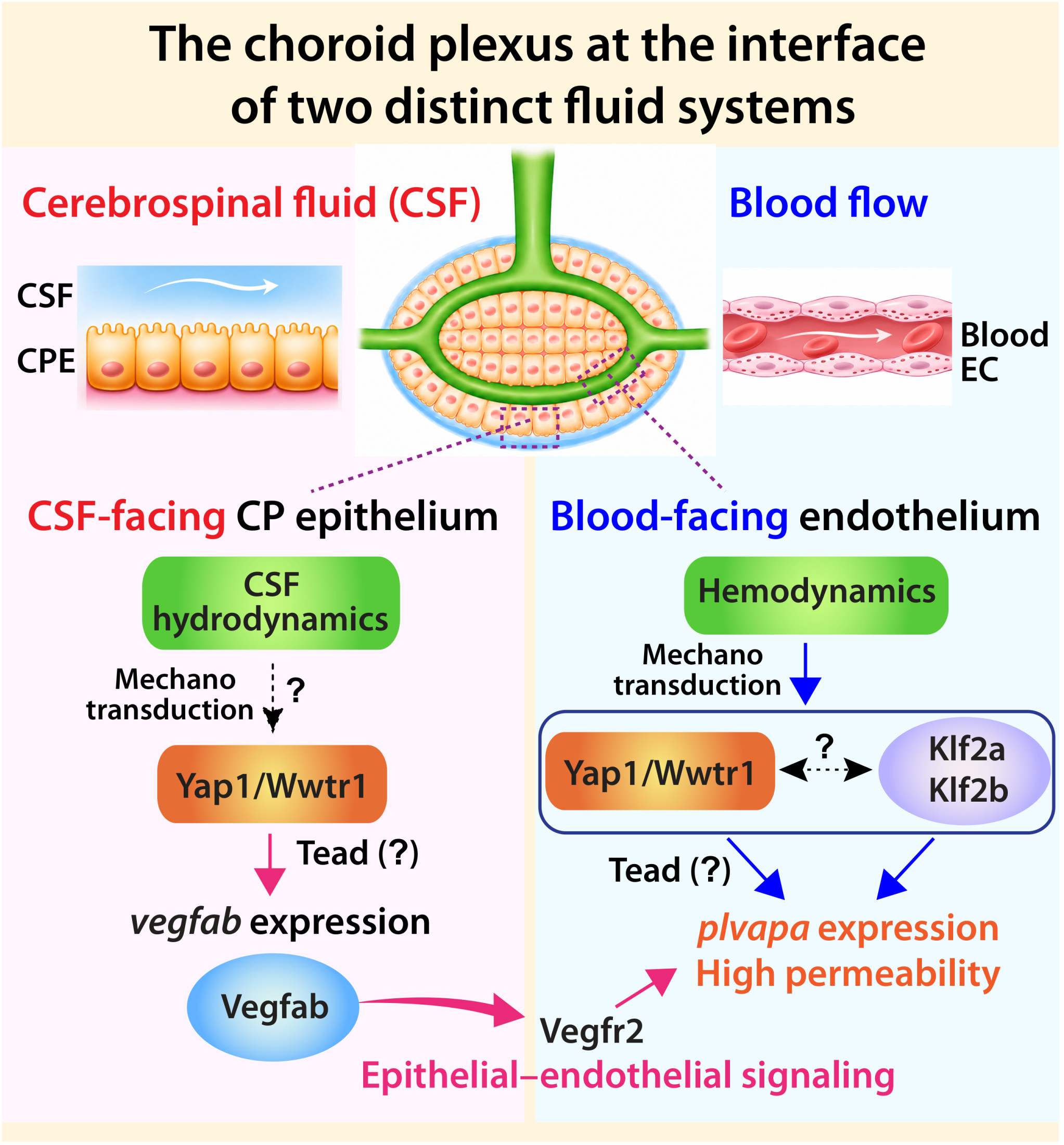
A model of epithelial-endothelial crosstalk for the establishment of EC identity and permeability in the mCP. Schematic diagrams illustrating the identified and proposed regulatory mechanisms governing fenestrated endothelial identity and vascular permeability in the mCP, which is uniquely positioned at the interface of two distinct fluid systems: blood and cerebrospinal fluid (CSF). In the CP epithelium, Yap1/Wwtr1 signaling regulates *vegfab* expression, which is crucial for high levels of *plvapa* expression in neighboring ECs, likely through epithelial-endothelial crosstalk. CSF hydrodynamics may contribute to epithelial Yap1/Wwtr1 activation through mechanotransduction. In the CP endothelium, Vegfr2 signaling and hemodynamics contribute to the regulation of *plvapa* expression and vascular permeability, with Klf2a/Klf2b and Yap1/Wwtr1 playing important roles in these processes. Yap1/Wwtr1 may control epithelial *vegfab* and endothelial *plvapa* expression via Tead-dependent transcriptional mechanisms. Together, these findings suggest a model in which CSF and blood fluid dynamics provide distinct mechanical cues linking Yap1/Wwtr1 to endothelial and epithelial gene regulatory programs that establish specialized mCP vascular identity and permeability.

Our successful characterization of Klf2a/Klf2b provides a proof of concept for the utility of our newly developed genetic screening platform in zebrafish, paving the way for future mechanistic investigations of other candidate genes identified in this study. Furthermore, integrating F0 genetic disruption and quantitative gene expression analysis with tracer dye-based permeability assays enables systematic molecular and functional characterization of genes involved in endothelial identity and permeability *in vivo*. Combined with the zebrafish model’s technical advantages, including rapid organogenesis and imaging accessibility, these screening platforms make it a powerful *in vivo* model for investigating embryonic cerebrovascular functional dynamics that are technically challenging to study in mammals and thereby defining previously understudied developmental mechanisms.

The identified cooperative functions of Klf2a/Klf2b in regulating endothelial *plvapa* expression and high vascular permeability in the mCP motivated us to investigate the broader role of hemodynamic and other mechanical forces in this process. By employing multiple strategies for manipulating hemodynamics, we showed that blood flow and circulating erythrocytes are required for endothelial high *plvapa* expression and permeability in the mCP. Given previous studies reporting the critical role of hemodynamics in developmental cerebrovascular remodeling (*61*) and both hemodynamic and Klf2a-dependent arterial specialization in the cerebrovasculature (*31*), Klf2a/Klf2b likely serve as crucial transcriptional mediators between hemodynamics and CP endothelial properties. Although the precise underlying mechanisms remain to be determined, our findings extend the roles of hemodynamic and Klf2-dependent signaling to the acquisition of permeable cerebrovascular identity and properties.

Previous studies reported important roles for endothelial YAP/TAZ signaling in developmental angiogenesis, vascular maturation, and homeostasis (*43, 44, 52*). Our findings suggest their potential dual functions in governing cerebrovascular regionalization. We showed that genetic and pharmacological disruptions of their activity reduced endothelial *plvapa* expression without affecting *plvapb* levels in the mCP, accompanied by reductions in vascular permeability. Conversely, Lats inhibition, previously shown to enhance Yap1/Wwtr1 activity (*48, 49*), increased both *plvapa* and *plvapb* expression in mCP ECs. However, these expression changes did not result in a significant increase in mCP vascular permeability. It is possible that WT mCP vessels already exhibit high baseline permeability, with modest increases in endothelial *plvapa* and *plvapb* expression not leading to readily detectable increases in permeability. Alternatively, this may reflect limited sensitivity of the dye leakage assay or a nonlinear relationship between *plvap* expression and vascular permeability.

Blood flow-dependent regulation of endothelial YAP1/WWTR1 activity is a conserved feature of zebrafish and mammalian vascular systems (*29, 52–54*). Our epistatic analyses showed that enhancing Yap1/Wwtr1 activity via Lats inhibition restored endothelial *plvapa* expression under erythrocyte depletion, indicating that Yap1/Wwtr1 signaling can counteract the effects of reduced hemodynamic input and supporting a role for Yap1/Wwtr1 downstream of hemodynamic signaling. In contrast, we found an unexpected interplay between Yap1/Wwtr1 and Vegfr2 signaling. Unlike its ability to restore *plvapa* expression under erythrocyte depletion, Lats inhibition failed to rescue *plvapa* expression during Vegfr2 inhibition. These contrasting findings suggest distinct regulatory relationships between Yap1/Wwtr1 and hemodynamic activity or Vegfr2 signaling in mCP vascular specialization.

VEGFA signaling promotes endothelial fenestration formation and PLVAP expression *in vitro* (*16, 17*) and maintains fenestrated vascular properties in adult tissues, including the CP (*14, 15*). Although VEGF signaling is a major regulator of vascular permeability, the upstream mechanisms governing its expression in specialized vascular environments remain poorly understood. Our findings reveal an upstream developmental mechanism that governs local epithelial Vegfa production required for the establishment of CP vascular identity and properties. The similar expression patterns of Yap1/Wwtr1 and *vegfab* in the mCP epithelium led us to investigate their regulatory relationship. By employing *yap1/wwtr1* loss-of-function and gain-of-function models, we found that Yap1/Wwtr1 activity positively controls epithelial *vegfab* expression. Given that *vegfab* functional loss and targeted ablation of Vegfab-expressing cells both reduced endothelial *plvapa* expression in the mCP, these findings together support a model in which Yap1/Wwtr1-dependent epithelial Vegfab production drives endothelial specialization and permeability through endothelial Vegfr2 signaling in the mCP. This model aligns with previous work reporting YAP/TAZ-dependent regulation of local Vegf signaling by intestinal stromal cells (*62*). Thus, Yap1/Wwtr1 appear to play distinct roles within mCP endothelial and epithelial cell populations, integrating mechanical and molecular signals to establish its unique vascular properties.

The presence of Yap1/Wwtr1 in both mCP epithelial and endothelial compartments raises the possibility that two distinct fluid environments contribute to mechanosensitive Yap1/Wwtr1 activity. In light of previous work highlighting the diverse roles of fluid mechanics in shaping vascular development (*31, 52, 61, 63, 64*), it will be particularly intriguing to investigate how systemic blood circulation and the CNS-specific CSF system intersect at the CP interface. We speculate that CSF hydrodynamics may provide additional biophysical cues influencing Yap1/Wwtr1 activity in the CP epithelium, given that CSF movements in zebrafish begin during early development (*65, 66*). Whether Yap1/Wwtr1 directly or indirectly regulates epithelial *vegfab* and endothelial *plvapa* expression remains an important next question. Our identification of consensus Tead-binding motifs in the promoter regions of both genes raises the possibility that Tead-dependent transcriptional mechanisms contribute to their regulation.

In summary, our study uncovers that blood flow-dependent physiological cues and an epithelial-endothelial biochemical signaling axis serve as developmental determinants of CP endothelial identity and permeability, thereby advancing our mechanistic understanding of cerebrovascular regionalization. Given the evolutionary conservation of CP anatomy, physiology, and molecular profiles (*60*), the developmental mechanisms identified here may have broad implications for CP vascular specialization across vertebrates. Since fenestrated vascular beds are present in diverse organs beyond the CNS, our findings may also provide broader insights into the molecular and physiological mechanisms underlying the establishment and functional specialization of fenestrated vasculature across different organs. Lastly, our integrated experimental approaches provide a technological foundation for further molecular and functional dissection of CP development and cerebrovascular diversity *in vivo*.

## Supporting information

Combined Supplemental Figures

## Abbreviations

ACeV: Anterior cerebral vein
BBB: Blood-brain barrier
CNS: Central nervous system
CP: Choroid plexus
CSF: Cerebrospinal fluid
CVOs: Circumventricular organs
dCP: Diencephalic choroid plexus
DLV: Dorsal longitudinal vein
EC: Endothelial cell
MCeV: Middle cerebral vein
mCP: Myelencephalic choroid plexus
MsV: Mesencephalic vein
PCeV: Posterior cerebral vein
PGV: Pineal gland vessel
PrA: Prosencephalic artery
Vegfs: Vascular endothelial growth factors
Vegfrs: Vascular endothelial growth factor receptors

## Acknowledgements

We thank Dr. Michael Taylor for providing the *Et(cp:EGFP)* line; Don Zeisloft and his team for zebrafish care and husbandry; Dr. Judith Drazba and her team for time-lapse and confocal imaging; and all members of the Matsuoka laboratory for discussions and comments on the manuscript. This work was supported by funding from the National Institutes of Health (R01 NS117510 and R21 NS142767) and start-up funds from the Cleveland Clinic Foundation to R.L.M.

## Author contributions

J.X.L. and N.J.L. performed CRISPR/Cas9-based F0 genetic screening by establishing the *plvapa*/*plvapb* RNAscope assay protocol and quantification methods and contributed to subsequent mechanistic studies. V.T. established the dye tracer permeability assay and quantification methods, performed all associated data quantification, and conducted spatial gene expression characterization. E.N.K. performed all *vegfab* mRNA quantifications in mCP epithelial cells. A.A.G. and F.Z.B. contributed to data collection related to phenotypic and expression analyses. R.L.M. designed and supervised the project, and wrote the manuscript. All authors have read and approved the final version of the manuscript.

## Competing interests

The authors declare that they have no competing interests.

## Data and materials availability

All data supporting the findings of this study are included within this manuscript. Correspondence and requests for materials should be addressed to Ryota L. Matsuoka.

## Materials and Methods

### Zebrafish husbandry and strains

All zebrafish husbandry was performed under standard conditions in accordance with institutional and national ethical and animal welfare guidelines. All zebrafish work was approved by the Cleveland Clinic Institutional Animal Care and Use Committee under protocol number 00003285. The following lines were used in this study: *Tg(kdrl:EGFP)^s843^* (*67*); *Tg(kdrl:NLS-mCherry)^is4^* (*68*); *Tg(kdrl:Has.HRAS-mcherry)^s896^*(*69*), abbreviated *Tg(kdrl:ras-mCherry)*; *Tg(gata1a:DsRed)^sd2^* (*70*); *Tg(hsp70l:sflt1, cryaa-cerulean)^bns80^* (*71*), abbreviated *Tg(hsp70l:sflt1)*; *Tg(hsp70l:sflt4, cryaa-cerulean)^bns82^* (*71*), abbreviated *Tg(hsp70l:sflt4)*; *TgBAC(vegfab:gal4ff)^bns273^* (*72*); *TgBAC(vegfd:gal4ff)^lri95^* (*21*); *Tg(UAS-E1b:NfsB-mCherry)^c264^* (*73*), abbreviated *Tg(UAS:NTR-mCherry)*; *Et(cp:EGFP)^sj2^* (*74*); and *vegfab^bns92^* (*75*). Adult fish were maintained on a standard 14 h light/10 h dark daily cycle. Embryos/larvae were raised at 28.5°C. To prevent skin pigmentation, 0.003% phenylthiourea (PTU) was added beginning at 10-12 hpf for imaging. Larvae analyzed at 10 dpf were transferred to tanks containing approximately 250 mL of water supplemented with 0.003% PTU (up to 25 larvae/tank) and fed with Larval AP100 (<50 μm dry diet, Zeigler) starting at 5 dpf.

### High-resolution melt analysis (HRMA)

A CFX96 Touch Real-Time PCR Detection System (Bio-Rad) was used for PCR reactions and subsequent HRMA. Precision melt supermix for high-resolution melt analysis (Bio-Rad) was used for all reactions. PCR conditions were as follows: 95°C for 2 min, 46 cycles of 95°C for 10 s, and 60°C for 30 s. Following the PCR, a high-resolution melt curve was generated by collecting EvaGreen fluorescence data over a temperature range of 65–95°C. Data were analyzed using normalized derivative plots. Genotyping of *vegfab^bns92^* mutant fish was performed by HRMA of PCR products as described previously (*21*).

### Generation of F0 knockouts (crispants)

Dual-guide ribonucleoprotein complexes (dgRNPs) consisting of crRNA, tracrRNA, and Cas9 protein were prepared as previously described (*26, 27*). All crRNAs employed for genetic screening were used after validating efficient disruption of their respective target loci, as previously described (*26, 27*). The specific sequences of all crRNAs used in this study are provided in Table S1.

For disruption of a single gene, three crRNAs per gene were co-injected with Cas9 protein into one-cell-stage embryos. Injection cocktails (4 μL) containing dgRNPs targeting each gene were prepared as follows: 0.4 μL each of 25 μM crRNA1:tracrRNA, crRNA2:tracrRNA, crRNA3:tracrRNA; 1.2 μL of 25 μM Cas9 protein; 1.2 μL of H2O; and 0.4 μL of 0.5% phenol red solution. For simultaneous disruption of two genes, six crRNAs targeting both genes (three crRNAs per gene) were co-injected with Cas9 protein into one-cell-stage embryos. Injection cocktails (4 μL) containing dgRNPs targeting two genes were prepared as follows: 0.4 μL each of 25 μM crRNA1:tracrRNA, crRNA2:tracrRNA, and crRNA3:tracrRNA targeting gene 1, and crRNA1:tracrRNA, crRNA2:tracrRNA, and crRNA3:tracrRNA targeting gene 2; 1.2 μL of 50 μM Cas9 protein; and 0.4 μL of 0.5% phenol red solution.

Injection cocktails were freshly prepared on the day of injection and were incubated at 37°C for 5 min and kept at room temperature prior to microinjection. Approximately 2 nL of the injection cocktails was injected into the cytoplasm of one-cell stage embryos. For phenotypic quantifications, larvae displaying obvious gross morphological defects were excluded from analyses, regardless of whether they were injected or uninjected.

### Immunohistochemistry

Immunohistochemistry was performed by following standard immunostaining procedures as described previously (*21*). Fish embryos/larvae were fixed in 4% paraformaldehyde (PFA) in phosphate-buffered saline (PBS), adjusted to pH 7.0, overnight at 4°C and dehydrated through immersion in methanol serial dilutions (50%, 75%, and 100%, followed by two additional washes in 100%, 10 min each) at room temperature (RT). Dehydrated samples were stored in 100% methanol at -20°C until use. Samples were rehydrated through immersion in methanol serial dilutions (50% and 25%, 10 min each) at RT and washed briefly three times in 1% PBST (1% Triton X-100 in 0.1M PBS) Samples were then permeabilized with 10 µg/ml Proteinase K in 1% PBST for 15 min at RT and blocked for 2-4 h at RT in blocking solution containing 0.5% bovine serum albumin in 1% PBST.

For immunohistochemistry using an anti-YAP/TAZ (D24E4) antibody, overnight PFA fixed samples were dehydrated in the same manner as described earlier. After rehydration through immersion in methanol serial dilutions (50%, 25%, then 1% PBST), samples were incubated with 3% H2O2 in water for 40 min at RT followed by 100% acetone treatment at 4°C for 8 min. The samples were briefly washed in 1% PBST twice at RT and then incubated in the blocking solution for 2 hr.

The following primary antibodies were used: chicken anti-GFP (GFP-1010; Aves Labs, 1:1000), rabbit anti-DsRed (#632496; Clontech, 1:300), and rabbit anti-YAP1/WWTR1 (D24E4) (#8418S; Cell Signaling Technology, 1:100). Following overnight incubation with primary antibodies at 4°C, samples were washed, incubated with appropriate secondary antibodies (1:500) overnight at 4°C, washed again, and processed for imaging as described previously (*21*).

### Pharmacological treatment

Larvae carrying either the *Tg(kdrl:EGFP)^s843^* or *Et(cp:EGFP)^sj2^* transgene were used for pharmacological experiments. For RNAscope assays, treatments were conducted either from 77 to 125 hpf for 48 h or from 101 to 125 hpf for 24 h, as indicated. For dye tracer vascular permeability assays, treatments were conducted from 72 to 96 hpf for 24 h prior to tracer injections. The following inhibitors were used: tricaine methanesulfonate (MS-222; Syndel), abbreviated tricaine, nifedipine (#N7634; Sigma), GsMTx4 (#4912; Tocris Bioscience), SKLB1002 (#SML1164; Sigma), MAZ51 (#676492; Sigma), verteporfin (#5305/10; Tocris Bioscience), and TDI-011536 (#E1314; Selleck Chemicals). Drug solutions were replaced with freshly prepared solutions every 24 h to maintain consistent drug activity. As previously described (*76*), pharmacological reagents other than tricaine and GsMTx4 were initially prepared as 100X stock solutions in 100% DMSO and diluted 1:100 in egg water immediately before use. Tricaine and GsMTx4 were dissolved in H₂O. The final working concentrations used were: 150 or 450 mg/L tricaine, 30 µM nifedipine, 300 nM GsMTx4, 3 µM SKLB1002, 30 µM MAZ51, 50 µM verteporfin, and 5 µM TDI-011536.

### Metronidazole (MTZ) treatment

For the cell ablation experiments in this study, MTZ substrate (Sigma, M3761) was used at 2 mM concentration dissolved in egg water containing 1% DMSO. Prior to treatment with 2 mM MTZ or 1% DMSO, embryos were manually dechorionated with forceps and incubated with freshly prepared 2 mM MTZ or 1% DMSO in egg water. Embryos were treated with 2 mM MTZ or 1% DMSO for indicated periods of time until sample fixation without replacing solutions.

### RNAscope *in situ* hybridization

RNAscope assays were performed on either *Tg(kdrl:EGFP)^s843^* or *Et(cp:EGFP)^sj2^* larvae collected at indicated developmental stages by following procedures described previously (*22*). Samples were fixed for two nights in 4% PFA/PBS, dehydrated, and rehydrated as described above for immunohistochemistry. Permeabilization was performed using Protease Plus (#322331; Advanced Cell Diagnostics, ACD) at 40°C for 3.5 min, 5 min, and 10 min for 75 hpf, 125 hpf, and 10 dpf samples, respectively, in the HybEZ^TM^ II hybridization system (#321711; ACD).

Gene-specific probes were designed and synthesized by ACD and prepared according to the manufacturer’s instructions. Probes against *plvapa* (#1330531-C3, 50X), *plvapb* (#1330521-C2, 50X), *klf2a* (#504421-C2, 50X), *vegfab* (#1896971-C2, 50X), and *kdrl* (#416611-C2, 50X) were freshly diluted to 1X in probe diluent (#300041; ACD) for each assay. Probes against *klf2b* (#574631, 1X) and *kdr* (#1318021-C1, 1X) were used without dilution. Most RNAscope assays were performed using two probes targeting different RNA transcripts, each assigned to a distinct detection channel (C1, C2, or C3). Probe hybridization was performed at 40°C for 2 h. During hybridization, 1X wash buffer was prepared from 50X stock (#310091; ACD) using H₂O and pre-warmed to 40°C.

Following hybridization, signal detection was performed using the RNAscope Multiplex Fluorescent Detection Kit v2 (#323110; ACD). Samples were washed three times in pre-warmed wash buffer at 40°C for 10 min each. Pre-warmed amplification reagents Amp 1, Amp 2, Amp 3 were then applied sequentially at 40°C for 30, 30, and 15 min, respectively, with two intervening washes in wash buffer at RT for 5 min each. Following Amp3 incubation, samples were washed twice in wash buffer at RT for 5 min each and kept overnight at RT in 5X SSC (750 mM NaCl, 75 mM sodium citrate, pH 7.0).

The following day, fluorescent signals were developed sequentially for each probe in dual-probe RNAscope assays (e.g., *plvapa/plvapb*, *klf2a/klf2b*, or *kdr/kdrl*). Samples kept in 5X SSC were washed once in wash buffer for 5 min at RT and incubated with the appropriate pre-warmed HRP-conjugated reagent (HRP-C1, HRP-C2, or HRP-C3) at 40°C for 15 min, followed by two washes in wash buffer at RT for 5 min each. The fluorescent signal for the first probe was developed using TSA Vivid Fluorophore 650 (#323273; ACD) diluted at 1:750 in Multiplex TSA buffer (#322810; ACD) at 40°C for 30 min. Following fluorescent signal development, samples were protected from light throughout the remaining steps of the assay until imaging to minimize photobleaching. After two washes in wash buffer at RT for 5 min each, samples were incubated with pre-warmed Multiplex FL v2 HRP Blocker (#323107; ACD) at 40°C for 15 min. After two washes in wash buffer at RT for 5 min each, samples were incubated with the appropriate pre-warmed HRP-conjugated reagent (HRP-C1, HRP-C2, or HRP-C3) at 40°C for 15 min, followed by two washes in wash buffer at RT for 5 min each. The fluorescent signal for the second probe was developed using TSA Vivid Fluorophore 570 (#323272; ACD) diluted at 1:750 in Multiplex TSA buffer at 40°C for 30 min. After two washes in wash buffer at RT for 5 min each, samples were incubated with pre-warmed Multiplex FL v2 HRP Blocker (#323107; ACD) at 40°C for 15 min. Following this step, samples were washed once in 1% PBST at RT and incubated with the anti-GFP antibody described above for at least 4 h at RT. After three washes in 1% PBST at RT for 20 min each, samples were incubated with Alexa Fluor Plus 488-conjugated goat anti-chicken IgY secondary antibody (#A32931; Thermo Fisher Scientific, 1:500) overnight at 4°C. After six washes in 1% PBST at RT for 15 min each, samples were processed for imaging.

### Quantification of *plvapa* and *plvapb* mRNA expression in mCP endothelial cells

Quantification was performed using confocal *z*-stack images of larvae carrying the *Tg(kdrl:EGFP)^s843^*reporter following wholemount RNAscope analysis of *plvapa* and *plvapb* expression. Given the spatial heterogeneity of transcript abundance within the mCP vasculature, four EC bodies along the outer boundaries of the mCP vascular loop, two on the rostral side and two on the caudal side, were selected for quantification in each larva, as illustrated in Fig. 1k. When EC bodies at these positions could not be clearly delineated or were judged unsuitable for reliable quantification due to their morphology, size, or angle, three EC bodies were quantified instead. The overall spatial distribution of mRNA signals in each sample was examined relative to that observed in WT larvae at the corresponding developmental stage, while ECs were selected based on consistent anatomical locations along the mCP vascular loop, independently of their transcript expression levels. To ensure rigorous and accurate quantitative comparisons, control and experimental larvae were processed simultaneously under the same RNAscope staining conditions whenever possible. Fluorescence signal scaling was adjusted to clearly distinguish individual mRNA puncta while avoiding excessive signal enhancement that obscured puncta boundaries or amplified weak background signals. Only discrete mRNA puncta with clearly discernible boundaries were counted. Individual optical sections of confocal *z*-stacks were examined, as needed, to distinguish overlapping RNA puncta and avoid duplicate counting across adjacent optical sections. For each transcript, the numbers of mRNA puncta associated with the selected individual EC nuclei were manually counted and averaged to obtain a single measurement of transcript abundance per EC nucleus for each larva. Each data point represents one larva.

### Quantification of *vegfab* mRNA expression in mCP epithelial cells

Quantification was performed using 125 hpf larvae carrying the *Et(cp:EGFP)^sj2^* reporter. *yap1* and *wwtr1* crispants were generated by injecting CRISPR RNA/Cas9 cocktails at the one-cell stage, as described earlier. Larvae were treated with 5 µM TDI-011536 for 48 h between 77 and 125 hpf. All samples were collected at 125 hpf. RNAscope assays using a *vegfab* probe were combined with anti-GFP immunostaining to simultaneously visualize *Et(cp:EGFP)* expression and *vegfab* transcripts. Following staining, high-magnification *z*-stack confocal images of the mCP region were captured at 1 µm intervals. A defined square area (20 µm × 20 µm) was placed near the center of the mCP to quantify and compare *vegfab* transcript abundance across experimental conditions. The number of mRNA puncta overlapping with *Et(cp:*EGFP)-positive mCP epithelial cells was manually counted by carefully examining individual *z*-planes of each image. Positive mRNA puncta were defined as those exhibiting signals brighter than the background fluorescence detected with the probe. In the same region, the number of epithelial cells was manually counted by examining cell boundaries between adjacent epithelial cells across individual *z*-planes to calculate *vegfab* transcript abundance per epithelial cell. The data quantified using this method well reflect our visual observations of gene expression and epithelial cell patterns.

### Heat shock treatments

Fish embryos/larvae raised at 28.5°C were subjected to a heat shock at 37°C for 1 h by replacing egg water with pre-warmed (37°C) egg water, as described previously (*71, 76*). After each heat shock, embryos/larvae were kept at room temperature for 10 min to cool down and then returned to 28.5°C. For heat shock experiments, *Tg(hsp70l:sflt1, cryaa-cerulean)^bns80^*, *Tg(hsp70l:sflt4, cryaa-cerulean)^bns82^*, and their sibling animals lacking the *cryaa:*cerulean eye marker, all carrying the *Tg(kdrl:EGFP)^s843^* reporter, were subjected to a heat shock at 70, 82, 94, 106, and 118 hpf prior to fixation at 125 hpf.

### Dye tracer injections

Tetramethylrhodamine-conjugated 40 kDa dextran (#D1842; Thermo Fisher Scientific) and Alexa Fluor 647-conjugated 38 kDa wheat germ agglutinin (WGA) (#W32466; Thermo Fisher Scientific) were used for dye tracer permeability assays. These tracers were each prepared in H2O at a concentration of 2 mg/mL, mixed in equal volumes, and approximately 2 nL of the mixture was injected into the common cardinal vein of *Tg(kdrl:EGFP)^s843^* larvae at the indicated developmental stages. For individual injections of Alexa Fluor 647-conjugated WGA, approximately 2 nL of a 2 mg/mL solution was used. Injected live larvae were imaged approximately 30 min to 1 h after injection.

### Quantification of fluorescent tracer extravasation

Tracer permeability quantification was performed in a similar manner as previously described (*77*). Specifically, embryos and larvae carrying the *Tg(kdrl:EGFP)^s843^* transgene were used to visualize blood vessels. Dye fluorescence intensity within the mCP vascular loops and the dorsal longitudinal vein (DLV) was measured using maximum-intensity projections of *z*-stack images in ImageJ. Since each injected larva displayed different levels of circulating tracer, fluorescence intensity within the mCP vascular loops was normalized to the circulating tracer intensity measured in the DLV of the same larva.

To measure circulating tracer fluorescence, a rectangular region of interest (ROI) was positioned within the width of the DLV by avoiding regions containing pigmentation. Samples were excluded when extravascular fluorescence adjacent to the DLV exceeded approximately two-thirds of the intravascular fluorescence or when circulating and extravascular fluorescence could not be clearly distinguished.

For mCP extravasation measurements, ROIs were manually delineated along the inner boundaries of vascular loops using the freehand selection tool in ImageJ, excluding the vessel walls. For larvae exhibiting a single vascular loop, the entire enclosed region was measured regardless of the spatial distribution of extravascular fluorescence. When larvae exhibited vascular branches that subdivided the central mCP vascular loop, up to three of the largest resulting loops were selected per larva for quantification. For these larvae, fluorescence intensities were combined using an area-weighted average to obtain a single leakage measurement per larva. When pigmentation partially obscured a loop, only the unobstructed region was measured. When pigmentation completely obscured the mCP vascular loops, these larvae were excluded from quantification. For T-shaped vascular configurations, ROIs were defined by the lateral boundaries of the nearest PCeV endothelial cell bodies beneath the DLV, corresponding to the expected location of the mCP vascular loop.

### Measurement of heartbeat rates

Heartbeat rates were measured in live larvae at 125 hpf. Individual larvae were transferred to a drop of their respective treatment solution in a glass-bottom Petri dish without anesthesia. The number of heartbeats was manually counted over a 15 sec period by observing cardiac contractions under brightfield or fluorescence microscopy. Larvae exhibiting gross morphological features that clearly differed from those of other larvae under the same experimental conditions were not used for quantification. The number of heartbeats counted over 15 sec was multiplied by four to calculate the heartbeat rate in beats per minute.

### Identification of putative Tead-binding motifs

Genomic sequences spanning 5 kb upstream of the transcription start sites of *plvapa* (NCBI reference sequence, NM_001114577.1) and *vegfab* (NCBI reference sequence, NM_001328597.2) were retrieved using the UCSC Genome Browser based on the zebrafish GRCz11/danRer11 genome assembly. These upstream sequences were examined for the presence of consensus Tead-binding motifs (GGAATG, TGGAATG, and AGGAATG) and their reverse complements (*58*).

### Confocal and stereo microscopy

Fluorescence imaging of live and fixed zebrafish samples was conducted using a Leica TCS SP8 confocal laser scanning microscope. Fish embryos and larvae were anesthetized with a low dose of tricaine, embedded in 1% low melt agarose in glass-bottom Petri dishes (MatTek), and imaged using a 25X water-immersion objective lens. Image acquisition and analysis were performed using LAS X software (Version 3.7.0.20979). Brightfield images of anesthetized fish were captured using a Nikon SMZ-18 stereomicroscope. Image acquisition and analysis for brightfield imaging were conducted using NIS-Elements BR Imaging software (Version 5.10.01).

### Statistical analysis

Statistical differences in mean values among multiple groups were determined using a one-way analysis of variance (ANOVA) followed by Dunnett’s multiple comparisons test against uninjected controls for CRISPR/Cas9-based genetic screening. For all other multi-group comparisons, one-way ANOVA followed by Tukey’s multiple comparisons test was used. Statistical differences in mean values between two groups were determined using a two-tailed Student’s *t*-test. Fisher’s exact test was applied to assess statistical differences in the penetrance of observed phenotypes. Statistical analyses were conducted using GraphPad Prism version 11.0.2. The criterion for statistical significance was set at *P* < 0.05. Error bars represent SEM.

## Notes

### Competing Interest Statement

The authors have declared no competing interest.

### Summary of Updates

The manuscript has been updated with improved clarity and refinements.

