## Supplementary material for "Hemodynamic and Yap1/Wwtr1-dependent epithelial-endothelial signaling establish fenestrated endothelial identity and permeability in the choroid plexus": Combined Supplemental Figures

### Figure S1

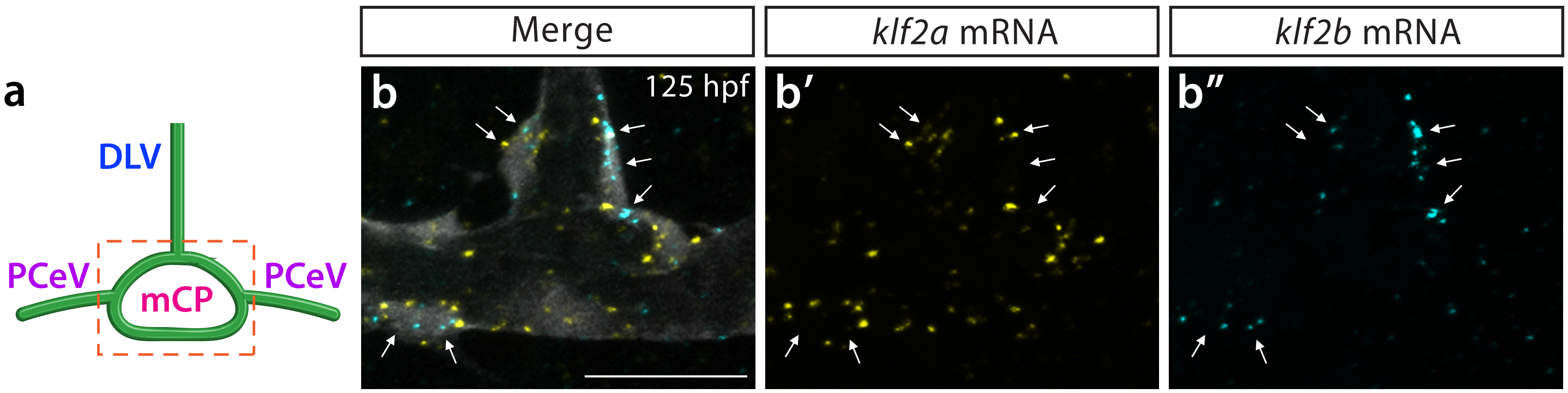


**Fig. S1. Endothelial *klf2a* and *klf2b* expression in the mCP at 125 hpf.**
(**a**) Schematic illustration of vessels contributing to the mCP vasculature (boxed area).
(**b-b”**) RNAscope analysis of *klf2a* (yellow, **b’**) and *klf2b* (cyan, **b”**) mRNA expression in the mCP vasculature visualized by the *Tg(kdrl:*EGFP*)* transgene at 125 hpf (merged image shown in **b**). Both *klf2a* and *klf2b* transcripts were detected within the same ECs (arrows) in this region.
Scale bars: 20 μm in **b** for **b-b”**.

Figure S2

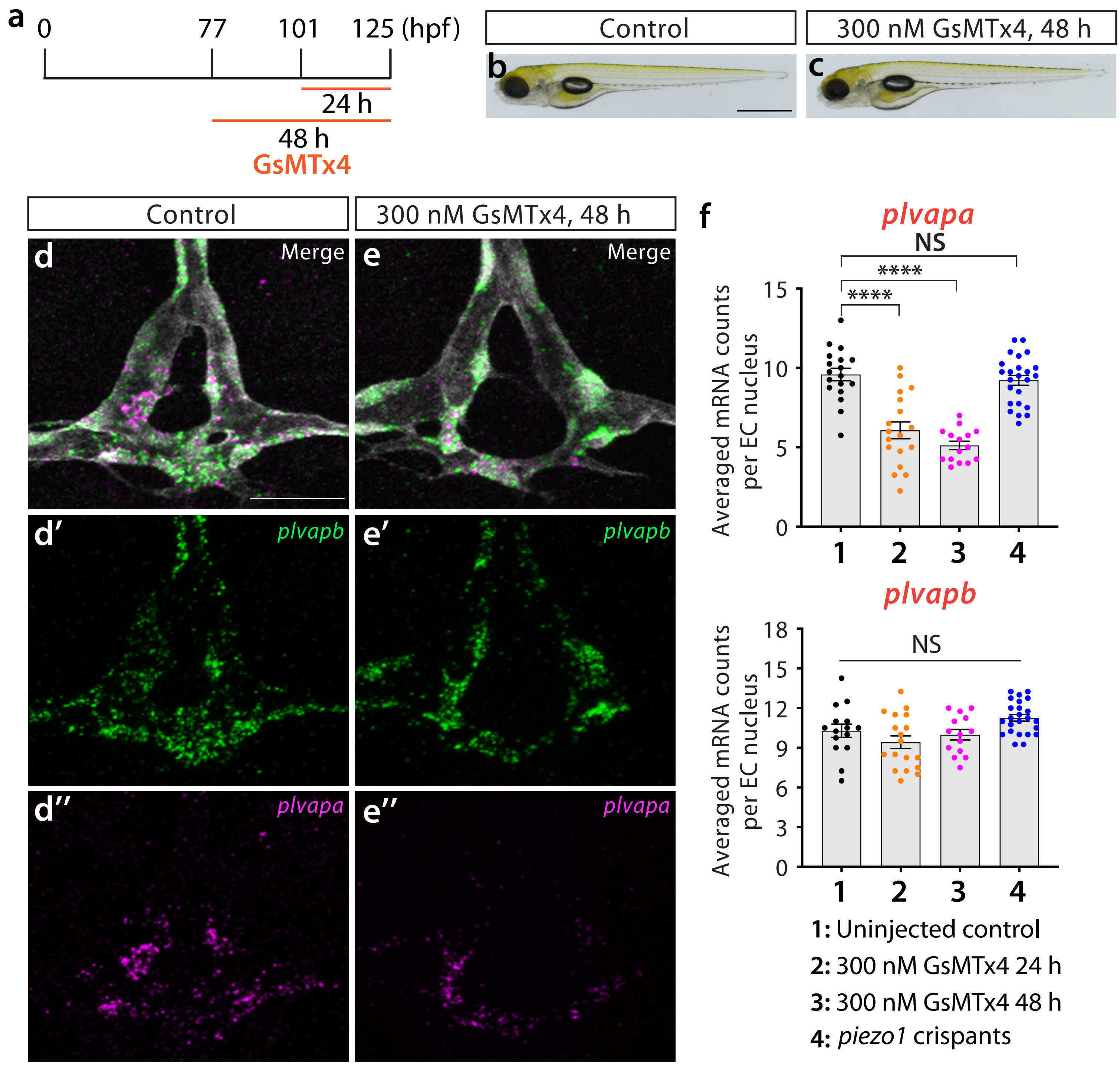


**Fig. S2. Mechanosensitive ion-channel activity contributes to mCP endothelial *plvapa* expression.**
(**a**) Experimental timeline for GsMTx4 treatment. Larvae were treated with 300 nM GsMTx4 for 24 h between 101 and 125 hpf or for 48 h between 77 and 125 hpf before sample collection at 125 hpf.
(**b**, **c**) Brightfield images of 125 hpf untreated control (**b**) and GsMTx4-treated larvae after 48 h (**c**).
(**d-e"**) Dorsal view images of *plvapb* (green, **d'**, **e'**) and *plvapa* (magenta, **d"**, **e"**) expression in the mCP of untreated control (**d-d"**) and GsMTx4-treated larvae after 48 h (**e-e"**) at 125 hpf. Merged images are shown in (**d**, **e**). GsMTx4 treatment reduced endothelial *plvapa* expression without affecting *plvapb* expression.
(**f**) Quantification of *plvapa* and *plvapb* mRNA abundance in mCP ECs of uninjected controls, larvae treated with 300 nM GsMTx4 for 24 or 48 h, and *piezo1* crispants at 125 hpf. GsMTx4 treatment reduced *plvapa* expression, whereas *piezo1* disruption produced no detectable *plvapa* reduction, suggesting that mechanosensitive ion-channel activity contributes to *plvapa* regulation through a mechanism that is not predominantly dependent on Piezo1. Across all conditions, *plvapb* expression remained largely unaffected.
Scale bars: 25 µm in **d** for **d-e"**; 1 mm in **b** for **b**, **c**.

### Figure S3

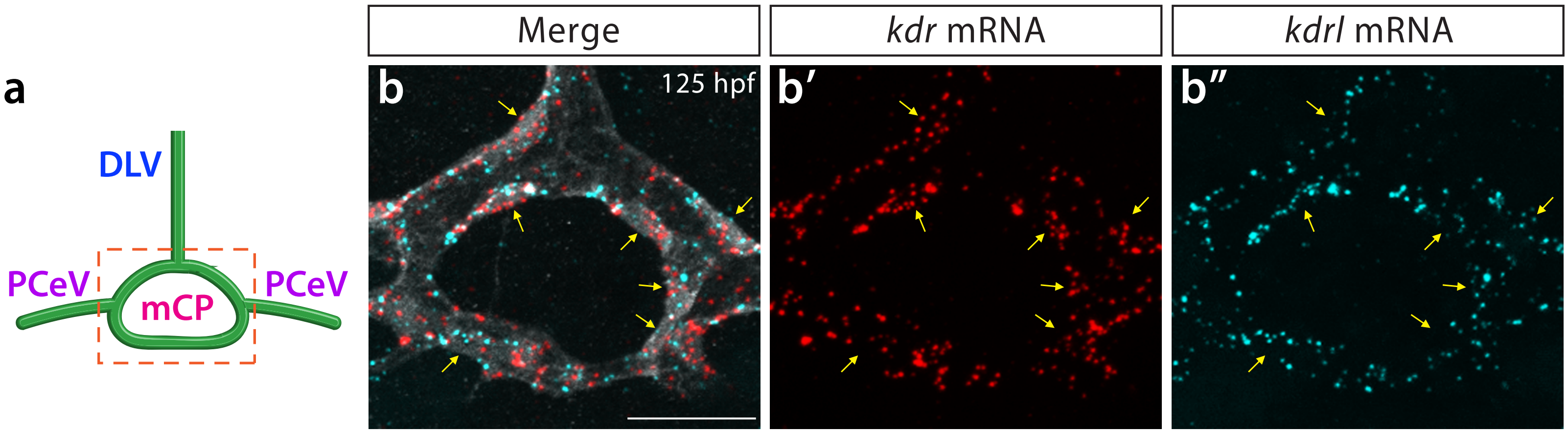


**Fig. S3. Endothelial *kdr* and *kdrl* expression in the mCP at 125 hpf.**
(**a**) Schematic illustration of vessels contributing to the mCP vasculature (boxed area).
(**b-b"**) RNAscope analysis of *kdr* (red, **b'**) and *kdrl* (cyan, **b"**) mRNA expression in the mCP vasculature visualized by the *Tg(kdrl:*EGFP*)* transgene at 125 hpf (merged image shown in **b**). Both *kdr* and *kdrl* transcripts were detected within the same ECs (arrows) in this region.
Scale bars: 20 µm in **b** for **b-b"**.

Figure S4

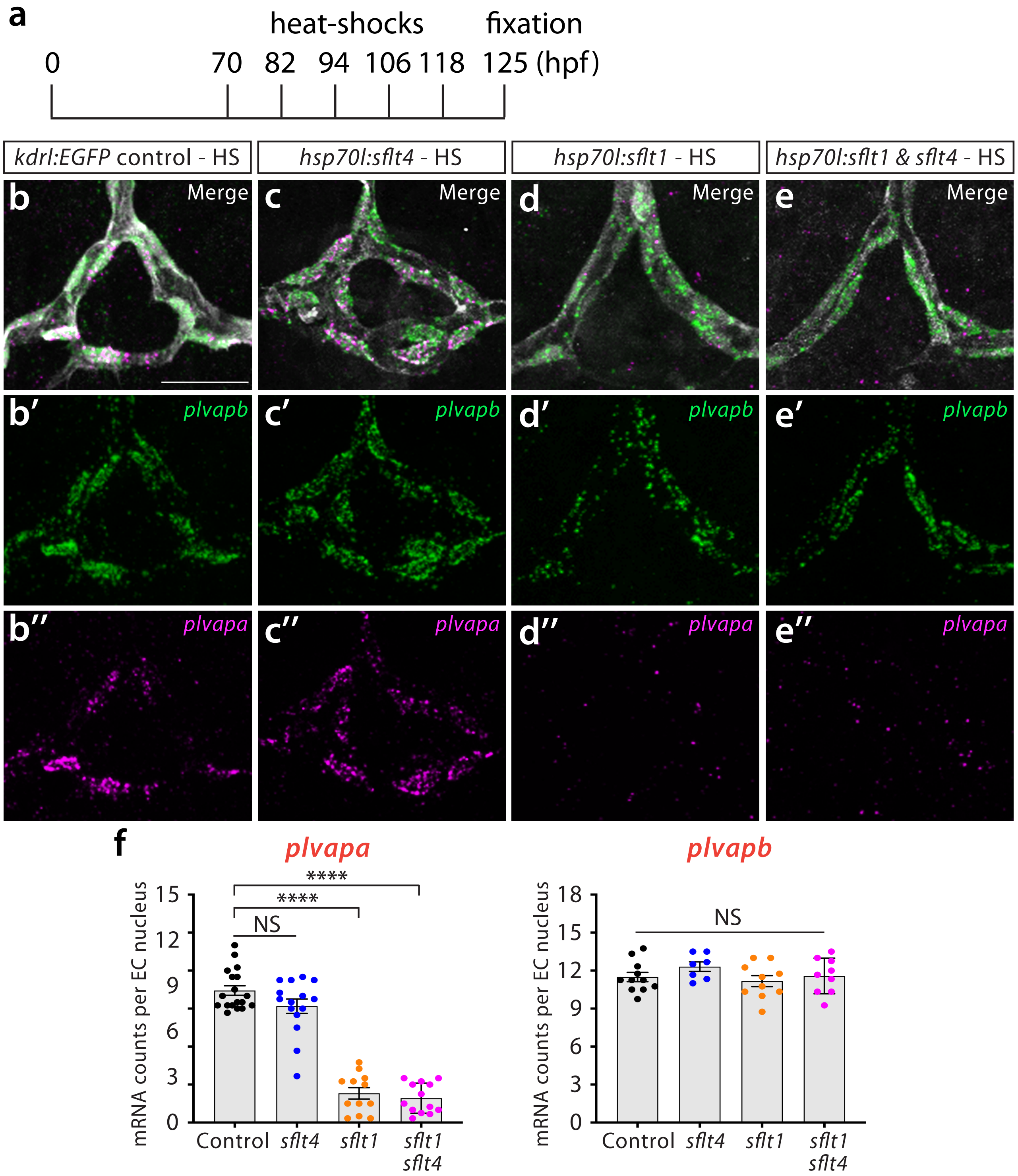


**Fig. S4. Inducible sequestration of Vegfa signaling selectively reduces endothelial *plvapa* expression in the mCP.**

(**a**) Experimental timeline for heat-shock-induced expression of soluble Flt4 (sFlt4) and/or soluble Flt1 (sFlt1). Larvae were subjected to repeated heat shocks from 70 to 118 hpf every 12 h and fixed at 125 hpf for analysis.

(**b-e"**) Dorsal view images of *plvapb* (green, **b'-e'**) and *plvapa* (magenta, **b"-e"**) expression in the mCP of heat-shocked larvae carrying *Tg(kdrl:EGFP)* (**b-b"**), *hsp70l:sflt4* (**c-c"**), *hsp70l:sflt1* (**d-d"**), or both *hsp70l:sflt1* and *hsp70l:sflt4* transgenes (**e-e"**) at 125 hpf. Merged images are shown in (**b-e**). Heat-shock-induced overexpression of sFlt1 drastically reduced endothelial *plvapa* expression without affecting *plvapb* expression, whereas sFlt4 overexpression did not alter the expression of either gene.

(**f**) Quantification of *plvapa* and *plvapb* mRNA abundance in mCP ECs under the indicated experimental conditions at 125 hpf.

Scale bars: 25 μm in **b** for **b-e"**.

### Figure S5

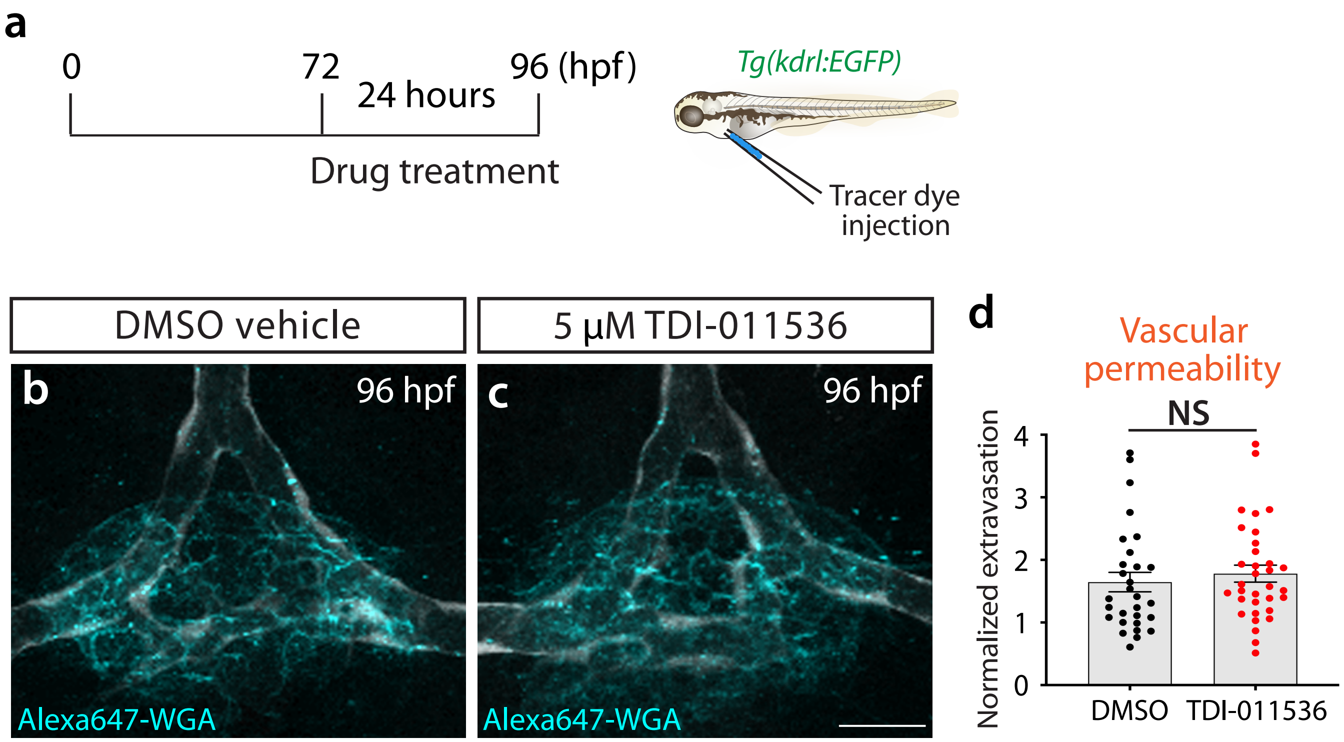


**Fig. S5. Pharmacological Lats inhibition does not significantly alter mCP vascular permeability.**
**(a) Experimental workflow for pharmacological Lats inhibition using the *Tg(kdrl:EGFP)* line. Larvae were treated with DMSO vehicle or 5 µM TDI-011536 for 24 h between 72 and 96 hpf and subsequently subjected to intravascular Alexa647-WGA tracer injection at 96 hpf.**
(**b**, **c**) Dorsal view images of Alexa Fluor 647-WGA extravasation in the mCP vasculature of DMSO-treated (**b**) and TDI-011536-treated (**c**) larvae at 96 hpf.
(**d**) Quantification of normalized WGA extravasation in the mCP vasculature of DMSO- and TDI-011536-treated larvae at 96 hpf, showing no significant difference in vascular permeability between the two groups (NS, not significant).
Scale bars: 20 µm in **c** for **b**, **c**.
