## Supplementary material for "Hemodynamic and Yap1/Wwtr1-dependent epithelial-endothelial signaling establish fenestrated endothelial identity and permeability in the choroid plexus": Table S1

| Gene | crRNA1 (5’ – 3') | crRNA2 (5’ – 3') | crRNA3 (5’ – 3') |
|---|---|---|---|
| *klf2a* | TCTTGACAAGTACCTGGATC | TGACAATCTCGGACTCGGCA | GTACAACCCGACTACATACT |
| *klf2b* | TCAGTGCCACGTTGAGCGGA | CAACACCACGGGCCCTCTGG | ATCTCCCACTCCGAGCCATG |
| *gata1a* | GGTTCATCTACATCAACGGG | ACGCTGGTGGAATAATACGA | TTCCCATCACGACTCGGGGA |
| *yap1* | ACCTCATCGGCACGGAAGGG | GAGTGTGCCCGGACGTGATG | GTCATCAGGTATCTCGTAAG |
| *wwtr1* | GATCCATGTCGCCAAAGACC | CGGACTCGGGCTCCCACTCC | GGTGTGGGAGTGGAGTCTCC |
| *piezo1* | CATCGGAGTGTGCACATGGT | GTATCAGAGCTCATTCGCCC | TGGCCAGTGTACGTCAATCC |
| *igfbp3* | CGCTCCTCGCAGCATTCGCG | CTGCGAGCCATGCGACGACG | GGCTGAAAGCGTCAGCCCTG |
| *irx3a* | CATCCGACCCATATACTCGC | TACGAGTTAAAGGACAGCCC | CCGAACTGGTACTGACCATA |
| *gata2a* | GAACACATCCACCTCGTCCG | ACCCCTACTACTCCAACTCT | TCACGGGAAGTCAGGTCTGT |
| *angpt1* | CAGCTACACGTTTATCCTCC | TGGTTGGACAAGTCGGCTTC | GAGCGCCGTCCACAACCACA |
| *esm1* | TTGCCCGGACAAATGCAACC | CGGACGGTGCTGGATGACTG | GACGCCATGCATCCCGGAGA |
| *cav1* | ACTTCAGGATGTCCACACCA | GCTTTGGGTCTCGGTTGACC | CCACACGCCGTCGAAGCTGT |
| *eng* | GCAACGACTGGATCGTCCTG | AAGTCCGTCCAGCATCCCAG | GGTTATCGAAAAGCATGGCG |
| *foxc1a* | CGGCACAACTCCCAGCGAGT | GCGTGAGAGTACATGGTCAT | AATGGCAATCCAGAACTCCC |
| *sox17* | GGTTTGGCTCGGATCGTCAC | TCAGTCATGATGCCTGGCAT | ACACGAGAAATGCTCTGCGG |
| *foxc1b* | CCTTATATCCCCGGAGACCA | CATACCGGTATATCCACCAC | ACGCACGAGCAGTACCCAGG |
| *aqp1a.1* | GTCTCCAAAGGGGACGCCCT | AGGTGTTGGAATTGAGCTCC | GAAGAGCAAGGCTTTCTGGC |
| *mef2d* | ACAACGAGCCACACGAAAGC | TTGGACACCGGCACTGTGAC | CTGGAGAAACCGTGTTCCTC |
| *ldb2a* | CCATGTACGGTGAATGTCTC | CATCTCGTAGATTCTGTACT | AGTAACGAGGGATAAGGGTT |
| *ripply3* | TGATCCGCGTCTACACACAC | TCTGTCGGGGAGAGACTCCA | GCGGACCATCAGCCCACTGC |
| *cby1* | GTCGACCCGTGAGATTGAAT | GGATCTCCTGTGATGAACAT | TCAAATTCGAAGACGGACAG |
| *mafba* | TGACATATTCCAACGCCAGC | AGGACGGCACGGAGCTGCAC | TGATCGGTGCCACGGCCCAC |
| *mafbb* | ACTACGGTGACTTTGACATG | TACTGGACCCTGAGCAGCAC | GAGGAAGGCACGGAGCTGCA |
| *TNFα* | AGGAGAGTTGCCTTTACCGC | CTTCAAAGTCGGGTGTATGG | TCCATCTTGTTGATTGCCCT |
| *hhex* | AAGACATCTTGGGAAGAACC | CGAGCCCACGCCGATACACC | CAAAGTCTCCCACGGAGCGC |
| *lztr1* | CCCACAAACTCATCGCACGG | TTGACGTGAAGGATTGCTCA | GAATATCTAACACTCAACTT |
| *tnfsf15* | GTGAGCGGTACTGTAACCGC | GACGCAGAATAGCACGCTGA | TGCGTGATACATAAGCGTGA |
| *sox18* | CCAGCACTCCGGGGCCTGAG | CAGAGCTGGCACCAGTGCCT | GAGCCCAGGGTCAAAGCTCC |
| *cyp26b1* | GAATGTGCGTAAGGTTCTTA | AGTGTCCATGGCACTTCTTC | GCTGAGGTGGACCGCAACAC |
